# Inhibition of CD226 Co-Stimulation Suppresses Diabetes Development in the NOD Mouse by Augmenting Tregs and Diminishing Effector T Cell Function

**DOI:** 10.1101/2024.07.16.603756

**Authors:** Matthew E. Brown, Puchong Thirawatananond, Leeana D. Peters, Elizabeth J. Kern, Sonali Vijay, Lindsey K. Sachs, Amanda L. Posgai, Maigan A. Brusko, Melanie R. Shapiro, Clayton E. Mathews, Rhonda Bacher, Todd M. Brusko

**Affiliations:** Diabetes Institute, College of Medicine, University of Florida, Gainesville, FL 32610; Department of Pathology, Immunology, and Laboratory Medicine, College of Medicine, University of Florida, Gainesville, FL 32610; Department of Biostatistics, College of Public Health and Health Professions, University of Florida, Gainesville, FL 32610; Department of Pediatrics, College of Medicine, University of Florida, Gainesville, FL 32610; Department of Biochemistry and Molecular Biology, College of Medicine, University of Florida, Gainesville, FL 32610

**Keywords:** CD226, disease prevention, monoclonal antibody, NOD mouse, type 1 diabetes

## Abstract

**Aims/hypothesis:** Immunotherapeutics targeting T cells are crucial for inhibiting autoimmune disease progression proximal to disease onset in type 1 diabetes. A growing number of T cell-directed therapeutics have demonstrated partial therapeutic efficacy, with anti-CD3 (α-CD3) representing the only regulatory agency-approved drug capable of slowing disease progression through a mechanism involving the induction of partial T cell exhaustion. There is an outstanding need to augment the durability and effectiveness of T cell targeting by directly restraining proinflammatory T helper type 1 (Th1) and type 1 cytotoxic CD8^+^ T cell (Tc1) subsets, while simultaneously augmenting regulatory T cell (Treg) activity. Here, we present a novel strategy for reducing diabetes incidence in the NOD mouse model using a blocking monoclonal antibody targeting the type 1 diabetes-risk associated T cell co-stimulatory receptor, CD226.

**Methods:** Female NOD mice were treated with anti-CD226 between 7-8 weeks of age and then monitored for diabetes incidence and therapeutic mechanism of action.

**Results:** Compared to isotype-treated controls, anti-CD226 treated NOD mice showed reduced insulitis severity at 12 weeks and decreased disease incidence at 30 weeks. Flow cytometric analysis performed five weeks post-treatment demonstrated reduced proliferation of CD4^+^ and CD8^+^ effector memory T cells in spleens of anti-CD226 treated mice. Phenotyping of pancreatic Tregs revealed increased CD25 expression and STAT5 phosphorylation following anti-CD226, with splenic Tregs displaying augmented suppression of CD4^+^ T cell responders *in vitro.* Anti-CD226 treated mice exhibited reduced frequencies of islet-specific glucose-6-phosphatase catalytic subunit related protein (IGRP)-reactive CD8^+^ T cells in the pancreas, using both *ex vivo* tetramer staining and single-cell T cell receptor sequencing (scTCR-seq) approaches. ^51^Cr-release assays demonstrated reduced cell-mediated lysis of beta-cells by anti-CD226-treated autoreactive cytotoxic T lymphocytes.

**Conclusions/interpretation:** CD226 blockade reduces T cell cytotoxicity and improves Treg function, representing a targeted and rational approach for restoring immune regulation in type 1 diabetes.

**Research in Context:** *What is already known about this subject?:*

- The co-stimulatory receptor CD226 is upregulated upon activation and is highly expressed on NK cell subsets, myeloid cells, and effector T cells.
- A single nucleotide polymorphism in CD226 (*rs763361*; C>T) results in a Gly307Ser missense mutation linked to genetic susceptibility for type 1 diabetes.
- Global knockout of *Cd226* and conditional *Cd226* knockout in FoxP3^+^ Tregs reduced insulitis severity and diabetes incidence in NOD mice, indicating a crucial role for CD226 in disease pathogenesis.

*What is the key question?:*

- Can CD226 blockade reduce T cell cytotoxicity and improve Treg function to diminish diabetes incidence in NOD mice?

*What are the new findings?:*

- Anti-CD226 treatment reduced insulitis, decreased disease incidence, and inhibited splenic CD4^+^ and CD8^+^ effector memory T cell proliferation.
- Pancreatic Tregs from anti-CD226 treated mice exhibited increased CD25 expression; splenic Tregs displayed augmented STAT5 phosphorylation and suppressive capacity *in vitro*.
- Anti-CD226 treatment reduced IGRP-specific pancreatic CD8^+^ T cell frequencies, and reduced autoreactive CD8^+^ T cell-mediated lysis of beta-cells *in vitro*.

*How might this impact on clinical practice in the foreseeable future?:*

- CD226 blockade could reduce autoreactive T cell cytotoxicity, enhance Treg function, and slow disease progression in high-risk or recent-onset type 1 diabetes cases.

## Introduction

The activation and proliferation of autoreactive T cells with specificity for pancreatic islet beta-cell antigens and modified peptide epitopes represent a hallmark histopathological feature of type 1 diabetes [1]. This lymphocytic infiltration is thought to drive the autoimmune destruction of these insulin-producing cells through both direct and indirect mechanisms [2]. Identifying strategies to suppress these autoreactive T cells is critical to establishing durable immune tolerance. Monoclonal antibodies (mAbs) represent a viable therapeutic option for treating type 1 diabetes. Teplizumab (anti-CD3) has recently received regulatory approval as the first disease-modifying therapy for stage 2 type 1 diabetes [3, 4], and mAbs have been used for the treatment of other autoimmune diseases, such as anti-CD20 for rheumatoid arthritis (RA) and anti-JAK1 for Crohn’s disease [5, 6].

The co-stimulatory receptor CD226 is expressed on several immune cell subsets, including natural killer (NK) cells, myeloid cells, and T cells, with the CD8^+^ T cell subset having the highest surface expression of CD226 [7, 8]. After binding its ligand CD155 and self-dimerization, CD226 contributes to immune activation through the phosphorylation of its immunoreceptor tyrosine tail (ITT)-like motif, which propagates Ras/MAPK signaling, leading to the transcription of pro-inflammatory genes, including *IFNG* and *IL17A* [9–11]. This signaling cascade in T cells is thought to promote the differentiation of the pro-inflammatory Th1 and Th17 subsets and impede differentiation into the anti-inflammatory Th2 and induced regulatory T cell (iTreg) subsets [12, 13].

Genetic susceptibility for several human autoimmune diseases, including type 1 diabetes, celiac disease, RA, and multiple sclerosis (MS), has been associated with a single nucleotide polymorphism (SNP) in *CD226* (*rs763361*; C>T), which results in a missense mutation leading to a Gly^307^ to Ser^307^ amino acid substitution [14]. This SNP is proximally located to two phosphorylation sites within the ITT-like motif (Tyr^322^ and Ser^329^), and Ser^307^ has been implicated in augmenting phosphorylation of downstream signaling elements such as ERK in human CD4^+^ T cells [15–18]. Importantly, *Cd226* is located within the *Idd21.1* risk locus (Chr. 18) of the NOD mouse and is orthologous to human *CD226* [19]; therefore, the NOD mouse is an ideal model to study the impact of blocking anti-CD226 mAbs on disease progression.

Previous work by our laboratory has shown that the global knockout (KO) of *Cd226* in the NOD mouse model reduced the severity of insulitis and the frequency of diabetes incidence, indicating a crucial role of CD226 in disease pathogenesis in this model [20]. The association of CD226 with autoimmunity has been further highlighted by Wang *et al.*, who identified that deletion of *Cd226* alleviated inflammation severity in an experimental allergic encephalomyelitis (EAE) mouse model of MS [21], as well as Zhang *et al.*, who demonstrated an anti-CD226 blocking mAb (clone LeoA1) reduced incidence of EAE [22]. Using adoptive transfer studies, we have previously reported that CD226-expressing CD4^+^ T cells substantially contribute to diabetes progression in the NOD [20], with a Treg-specific conditional KO of *Cd226* demonstrating that CD226-expressing Tregs more readily destabilize into a pro-inflammatory “ex-Treg” phenotype, further driving diabetes progression [23]. Thus, we investigated whether employing a blocking mAb targeting CD226 to selectively inhibit inflammatory T cell activation could reduce autoreactivity.

Using the NOD mouse model of type 1 diabetes, we conducted an *in vivo* intervention study and *ex vivo* and *in vitro* mechanistic studies of an anti-CD226 blocking mAb. We investigated the differential impacts of CD226 blockade on conventional CD4^+^ T cell (Tconv), Treg, and CD8^+^ T cell compartments and how those effects may contribute to protection from disease to inform future translational efforts in human trials.

## Materials & Methods

### Animals

Female NOD/ShiLtJ (NOD; RRID: IMSR_JAX:001976) mice were purchased from The Jackson Laboratory (Bar Harbor, ME, USA) for *in vivo* intervention studies and *ex vivo* mechanistic studies. Female NOD/ShiLt-Tg(*Foxp3*-EGFP/cre)1cJbs/J (NOD-*Foxp3*^EGFP^; RRID: IMSR_JAX:008694) transgenic mice were used for Treg suppression studies [24]. Chromium-release assays were accomplished using cells from NOD/ShiLtJ-*Tcra*-AI4.*Tcrb*-AI4.*Thy1a*.*Rag1*^-/-^ mice (NOD-AI4; [25]). Animals treated with mAbs were housed in animal biosafety level 2 (aBSL-2) containment, whereas all others were housed in animal biosafety level 1 (aBSL-1) containment. All animals were housed in specific pathogen-free facilities at the University of Florida, with food and water available *ad libitum*. All studies were conducted in accordance with protocols approved by the University of Florida Institutional Animal Care and Use Committee (UF IACUC) and in accordance with the *Guide for Care and Use of Laboratory Animals* [26].

### Preparation of Single-Cell Suspensions

Isolation of cells from relevant tissues, including the spleen, thymus, pancreatic-draining lymph nodes (pLN), and pancreas, was necessary for *in vitro* and *ex vivo* assessment of the mAb mechanism of action. For all tissues except the pancreas, homogenous single-cell suspensions were generated by mechanically dissociating organs and filtering through a 70 µm membrane filter. Cell suspensions were washed with complete RPMI [cRPMI; RPMI 1640 media Phenol Red with L-Glutamine 139 (Lonza, Basel, CH-BS, Switzerland), 5mM HEPES (Gibco, Waltham, MA, USA), 5 mM MEM 140 Non-Essential Amino Acids (NEAA; Gibco), 2mM Glutamax (Gibco), 50 µg/mL penicillin 141 (Gibco), 50 µg/mL streptomycin (Gibco), 20 mM sodium pyruvate (Gibco), 50 mM 2-mercaptoethanol (Sigma-Aldrich, St. Louis, MO, USA), 20 mM sodium hydroxide (Sigma-Aldrich) and 10% FBS (Genesee Scientific, El Cajon, CA, USA)] and pelleted by centrifugation (350 x g for 7 min). Pancreas tissues were minced into 1 mm pieces and incubated in cRPMI media with 1 mg/mL collagenase IV (Gibco) for 18 minutes at 37°C. After digestion, pancreas suspensions were washed with cRPMI and strained through a 40 μm filter. Lysis of red blood cells (RBC) was accomplished by resuspension of cell pellets in ACK Lysing Buffer (Gibco) for 5 minutes at 4°C before quenching in 1x Phosphate Buffered Saline (PBS; Gibco). Cell viability was quantified by staining with Acridine Orange/Propidium Iodide at a 1:1 dilution in a Cellometer slide before reading on an Auto2000 Cellometer (Nexcelom Biosciences, Lawrence, MA, USA).

### Administration of Anti-CD226 and Isotype Control mAbs

Blocking mAb against CD226 (BioLegend, San Diego, CA, USA; clone 480.1, RRID: AB_2876467) and its corresponding rat IgG2a isotype control (BioLegend; RRID: AB_11147167) were obtained in a low-endotoxin, azide-free formulation. mAbs were diluted in PBS (Gibco) to 1.33 µg/µL for intraperitoneal (i.p.) administration. Mice received three 200 µg doses of mAb at 49, 53, and 56 days of age and were monitored daily for potential reactions.

### Anti-CD226 mAb Blockade Validation

For *in vitro* validation, NOD-*Foxp3*^EGFP^ splenocytes were resuspended at 0.5 x 10^6^ cells/mL and incubated with 20 µg/mL IgG2a isotype control (BioLegend) or anti-CD226 mAb (BioLegend) for 30 minutes in cRPMI at 37°C. Cells were stained with Live/Dead™ Near-IR viability dye (Invitrogen, Waltham, MA, USA) per the manufacturer’s protocol. Then, Fc receptors were blocked with 11-CD16/CD32 (BD Pharmingen, Franklin Lakes, NJ, USA; RRID: AB_394656) for 5 minutes at 4°C to prevent non-specific binding before extracellular staining with an antibody cocktail consisting of anti-mouse CD4-PerCP-Cy5.5 and CD811-BV711, and a fluorophore-conjugated anti-CD226 mAb (clone TX42.1) for 30 min at 23°C (clone, RRID, concentration, and manufacturer information provided in **Table S1**). Data were collected on an Aurora 5L (16UV-16V-14B-10YG-8R) spectral flow cytometer (Cytek, Freemont, CA, USA) and analyzed using FlowJo software (TreeStar; version 10.6.1).

### T Cell Proliferation Assays

Whole splenocytes isolated from 8- to 12-week-old female NOD mice were labeled with 5 µM CellTrace™ Violet (CTV; Thermo Fisher, Waltham, MA, USA) as recommended by the manufacturer’s protocol. Following proliferation dye staining, cells were resuspended at 0.5 x 10^6^ cells/mL and incubated with 20 µg/mL IgG2a isotype control (BioLegend) or anti-CD226 mAb (BioLegend) for 30 minutes in cRPMI at 37°C. 0.25 x 10^6^ cells were stimulated with either plate-bound 11-CD3 (BioLegend; RRID: AB_11149115; 2 µg/mL) and plate-bound 11-CD28 (BioLegend; RRID: AB_11147170; 1 µg/mL) or plate-bound 11-CD3 (BioLegend; 2 µg/mL) and plate-bound CD155-Fc (BioLegend; 1 µg/mL). Following 96 hours of culture, supernatants were stored at -20°C for ELISA described below, and cells underwent viability staining and 11-CD16/CD32 blocking, as described above, followed by surface staining with anti-mouse CD4-PerCP-Cy5.5 and CD811-BV711 for 30 min at 23°C (**Table S1**). Data were collected on a Cytek™ Aurora 5L spectral flow cytometer. The detailed gating strategy is shown in **Figure S1.** The proliferation of CD4^+^ and CD8^+^ T cells was established by the division index (DI) method using proliferation modeling on FlowJo software (TreeStar; version 10.6.1).

### ELISA

To determine whether anti-CD226 mAb blockade modulates secretion of the pro-inflammatory IFN-γ or anti-inflammatory IL-10 cytokines, ELISAs were performed on culture supernatants from the *in vitro* T cell proliferation assay described above. Briefly, culture supernatants were diluted 1:2 for IL-10 and 1:100 for IFN-γ, and ELISAs were performed using the Mouse IL-10 and Mouse IFN-γ OptEIA Kits (BD Biosciences) according to the manufacturer’s protocol. Colorimetric analyses were performed in duplicate at 450 nm with a λ correction at 570 nm on a SpectraMax M5 microplate reader (Molecular Devices, San Jose, CA, USA).

### Flow Cytometry

1-2 x 10^6^ cells from each tissue were used for flow cytometry and were stained with Live/Dead™ Near-IR viability dye. Cells were blocked with 11-CD16/CD32 (BD Pharmingen) and Brilliant Stain Buffer (BD Biosciences) for 5 minutes at 4°C before extracellular staining with an antibody cocktail consisting of anti-mouse CD4-PerCP-Cy5.5, CD8-BV711, CD25-Alexa Fluor 700, CD44-PE, CD62L-APC, and CD226-BV650 for 30 minutes at 23°C (**Table S1**). Next, cells were fixed and permeabilized using the eBioScience™ FOXP3 Transcription Factor Staining Buffer Set (Invitrogen) according to the manufacturer’s instructions. Permeabilized cells were stained with anti-mouse Foxp3-Alexa Fluor 488, Helios-Pacific Blue, and Ki-67-PE-Cy-7 antibodies overnight at 4°C (**Table S1**). Data were collected on a Cytek™ Aurora 5L spectral flow cytometer and analyzed using FlowJo software (TreeStar; version 10.6.1). Gating strategies were determined using fluorescence-minus one (FMO) and unstained controls. Detailed gating strategies for Tregs and T cell memory subsets are shown in **Figure S2**.

### Anti-CD226 mAb Blockade Persistence

To assess the *in vivo* persistence of anti-CD226 mAb, 1-2 x 10^6^ splenocytes underwent viability staining and 11-CD16/CD32 blocking, as previously described, before surface staining with anti-mouse CD4-PerCP-Cy5.5, CD8-BV711, NKp46-Alexa Fluor 647, and anti-rat IgG2a-FITC for 30 minutes at 23°C (**Table S1**). Flow cytometry data were collected and analyzed as described above. The gating strategy to assess anti-CD226 mAb persistence is shown in **Figure S3**.

### Histology

Pancreata were collected from 12-week-old mice at necropsy and fixed overnight in a buffered 10% formalin solution. Samples underwent paraffin-embedding, and three sections (250 µm steps) were obtained for H&E staining. Digital whole-slide scans of pancreas sections were obtained using an Aperio CS Scanner (Leica Biosystems, Wetzlar, Germany). Two blinded observers completed insulitis scoring for at least 45 islets per mouse (with one islet defined as >10 endocrine cells) according to previously published guidelines [27].

### Intervention Study

Beginning at seven weeks of age, concurrent with the initiation of mAb administration, blood glucose levels of female NOD mice were monitored weekly until 30 weeks of age, using an AlphaTrak™ glucometer (Zoetis, Parsippany, NJ, USA) to measure samples obtained from a tail vein bleed. Mice with blood glucose ≥ 250 mg/dL were retested the following day, and those with two consecutive blood glucose levels ≥ 250 mg/dL were diagnosed with diabetes. Body mass measurements were recorded weekly to monitor for possible impacts of treatment on the growth and overall health of the animals. Mice were humanely euthanized by CO_2_ asphyxiation and cervical dislocation at diabetes onset or study conclusion.

### pSTAT5 Phosflow Assay

To determine whether mAb blockade alters the JAK2-STAT5 pathway in Tregs, we quantified phosphorylated STAT5 (pSTAT5) levels following stimulation with recombinant human IL-2 (rhIL-2; Roche, Basel, CH-BS, Switzerland; [28]). Single-cell suspensions of splenocytes were obtained from 12-week-old NOD mice five weeks after *in vivo* treatment with IgG2a isotype control or anti-CD226 mAb, as described above. Cells were plated at 1 x 10^6^ cells/mL cRPMI. They were stimulated with 10 IU/mL of rhIL-2 for 0, 15, or 60 minutes before immediately fixing cells with warmed CytoFix Buffer (BD Biosciences) for 10 minutes at 37°C. Following fixation, cells underwent viability staining, as described above, before permeabilizing cells for 30 minutes at 4°C with chilled Perm Buffer III (BD Biosciences). Following Fc blocking with anti-CD16/32, cells were stained with anti-mouse CD4-PE-Cy7, CD8-BV711, Foxp3-AF488, and pSTAT5-AF647 overnight at 4°C (**Table S1**). Flow cytometry data were collected and analyzed as described above.

### Suppression Assays

To assess the potential impact of anti-CD226 mAb blockade on the suppressive capacity of mouse Tregs, we conducted *in vitro* suppression assays [29] using serial dilutions of fresh splenic Tregs with irradiated autologous whole splenocytes and autologous CD4^+^ Tconv that respectively served as antigen-presenting cells (APCs) and responder T cells (Tresp). Briefly, whole splenocytes were isolated from 8- to 12-week-old female NOD-*Foxp3*^EGFP^ mice. The splenocyte suspension was split so that half of the cells received ^137^Cs gamma irradiation at a dosage of 3000 cGray, after which cells were concentrated at 1.0 x 10^6^ cells/mL cRPMI, and 50,000 cells were added to each well of a 96-well U-bottom plate to provide APC-mediated stimulation alongside soluble 11-CD3 (BioLegend; 0.5 µg/mL) and 11-CD28 (BioLegend; 0.5 µg/mL).

Concurrently, Treg and Tconv cells were isolated from the remaining splenocytes by enriching for CD4^+^ T cells using the EasySep™ Mouse CD4^+^ T Cell Isolation Kit (StemCell, Vancouver, BC, Canada) according to the manufacturer’s instructions. CD4^+^ T cell-enriched splenocytes were stained with anti-CD4-PE-Cy7 (**Table S1**) to allow for both Foxp3^+^ Treg and Foxp3^-^ Tconv isolation by sorting for CD4^+^GFP^+^ and CD4^+^GFP^-^ cells, respectively, using a FACSMelody™ Cell Sorter (Becton Dickinson, Franklin Lakes, NJ, USA; **Figure S4**). Following isolation, Tconv were labeled with CTV, as described above, concentrated at 1 x 10^6^ cells/mL cRPMI, and 50,000 cells were added to each well. Before co-culture, Tregs were split and incubated at a concentration of 1 x 10^6^ cells/mL cRPMI for 30 minutes at 37°C in cRPMI with 10 µg/mL anti-CD226 mAb (BioLegend) or IgG2a isotype control (BioLegend). After incubation, cells were washed with cRPMI to remove any unbound mAb, then co-cultured with the autologous Tconv at the following Treg:Tresp ratios: 1:1, 1:2, 1:4, 1:8, 0:1. Following 72 hours of co-culture, cells underwent viability dye staining, Fc blocking with anti-CD16/32, surface staining with anti-mouse CD4-PerCP-Cy5.5, and flow cytometry data collection as previously described (**Table S1**). The detailed gating strategy is shown in **Figure S5**. Percent suppression of CD4^+^GFP^-^ Tresp was established by the Division Index (DI) method using proliferation modeling on FlowJo (TreeStar; version 10.6.1).

### IGRP_206-214_ Tetramer Staining

Single-cell suspensions obtained from collagenase-digested pancreas from female NOD mice were assessed for frequency of islet-specific glucose-6-phosphatase catalytic subunit-related protein (IGRP)_206-214_ reactive CD8^+^ T cells, five weeks following treatment with IgG2a isotype control or anti-CD226 mAb *in vivo*. Cells underwent viability dye staining and Fc blocking before concurrent surface antibody (anti-mouse CD4-PerCP-Cy5.5 and CD8-BV711; **Table S1**) and tetramer staining (10 nM IGRP_206-214_-BUV395 tetramer) for 1 hour at 37°C in the presence of 75 nM dasatinib [30]. Flow cytometry data were collected and analyzed as described above with a detailed gating strategy shown in **Figure S6**.

## Chromium-Release Assay

To evaluate whether anti-CD226 mAb blockade could sufficiently reduce the cytotoxicity of autoreactive CD8^+^ T cells and subsequently decrease pancreatic beta-cell killing, chromium-release assays were performed in the format previously described by Chen et al. [31]. Briefly, whole splenocytes isolated from 3- to 4-week-old NOD-AI4 mice, reported to possess diabetogenic insulin-reactive CD8^+^ T cells [32], were concentrated at 5 x 10^6^ cells/mL in cRPMI and incubated with either IgG2a isotype control or anti-CD226 mAb (BioLegend; 20 µg/mL) for 30 minutes at 37°C. After incubation, splenocytes were diluted to a concentration of 2 x 10^6^ cells/mL cRPMI and activated over three days in the presence of 0.1 µM AI4 mimotope (YFIENYLEL; GenScript, Piscataway, NJ, USA) and Teceleukin rhIL-2 at a concentration of 25 IU/mL cRPMI to selectively expand CD8^+^ T cells. Following activation, autoreactive CD8^+^ T cells were transferred into complete DMEM media (cDMEM; Dulbecco’s Modification of Eagle’s Medium (DMEM; Lonza), 5 mM HEPES (Gibco), 5 mM MEM NEAA (Gibco), 50 µg/mL penicillin 141 (Gibco), 50 µg/mL streptomycin (Gibco), 0.02% Bovine Serum Albumin (BSA; Sigma-Aldrich), and 10% FBS (Genesee Scientific)) and were added to flat-bottom 96 wells containing pre-seeded ^51^CrNa_2_O_4_-labeled (1.8 x 10^5^ Bq/well; Revvity, Waltham, MA, USA) murine NIT-1 pancreatic beta-cells (RRID: CVCL_3561; [33]) at the following effector:target cell ratios: 0:1, 1:1, 2:1, 5:1, 10:1, 25:1. Following 16 hours of co-culture, supernatants were removed and transferred to 6x50 mm lime glass tubes. Lysates of adherent cells were collected using a 2% SDsS wash and transferred into separate tubes. ^51^Cr activity, measured in counts per minute (CPM), was assessed for both fractions on a Wizard 1470 automatic gamma counter (Revvity). The specific lysis of NIT-1 cells was calculated as follows:

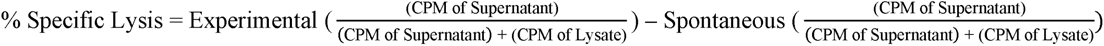

### Single Cell Sequencing

Single-cell suspensions from the pancreas and pLN were obtained from 12-week-old female NOD mice following treatment with IgG2a isotype control (pLN: n=3; pancreas: n=3) or anti-CD226 mAb (pLN: n=5; pancreas: n=2) to prepare sequencing libraries on the 10x Genomics platform. CD3^+^ T cells were enriched within each sample using the Mouse T cell EasySep™ kit (StemCell) per the manufacturer’s instructions. Each sample underwent Fc blocking with anti-CD16/32 for 10 minutes at 4°C before staining with oligonucleotide-tagged antibodies for CD4, CD811, CD44, CD62L, and TIGIT (clone and DNA barcode information provided in **Table S2**) for 30 minutes at 4°C. Following staining, cells were washed three times with PBS + 1.0% BSA before loading 5,000 CD3^+^ enriched cells onto a Chromium Next GEM Chip K (10x Genomics, Pleasanton, CA, USA) to generate Gel Beads in-Emulsions (GEMs) using a 10x Chromium Controller (10x Genomics). Gene expression, V(D)J, and feature barcode libraries were generated using the Chromium Next GEM Single Cell 5’ v2 and Chromium Single Cell Mouse T Cell Receptor (TCR) Amplification Kits (10x Genomics). All libraries were sequenced using the NovaSeq XPlus Illumina platform with a minimum sequencing depth of >19,214 reads/cell for gene expression (GEX) libraries, >5,271 reads/cell for V(D)J libraries, and >6,053 reads/cell for feature barcode (FB) libraries.

### Pre-processing of 10x Genomics Sequencing Data

The Cell Ranger (version 3.0.0, 10x Genomics) multi-pipeline was used to generate raw feature-barcode matrices by processing raw sequencing reads from GEX and FB libraries as well as annotated full-length transcripts (contigs) by processing the raw sequencing reads of V(D)J libraries. Briefly, GEX sequencing reads were aligned to a reference genome (mm10-2020-A) using STAR [34]. Confidently mapped reads sharing the same unique molecular identified (UMI), 10x barcode, and feature were collapsed, with the number of reads per feature saved in the raw FB matrix. Contigs with V(D)J segment labels were aligned to a reference genome (GRCm38-alts-ensembl-7.0.0) to identify complementarity-determining region 3 (CDR3) sequences. Productive contigs sharing the same UMI and barcode were saved as filtered contig annotations.

### Quality Control of scRNA-seq and CITE-Seq Data

Raw FB matrices of cells sequenced from each tissue were imported into R (version 4.3.1) using the Read10X function in the Seurat package (version 5.0.1). Live cell-containing droplets were distinguished using gene (80 – 2,000) and protein library size (1.0 – 3.0) as well as mitochondrial content (<15%). CITE-seq data were normalized using the denoised and scaled by background (dsb) method with the DSBNormalizeProtein function, without isotype controls, in the dsb package (version 1.0.3). To remove variation in our single cell RNA sequencing (scRNA-seq) depth between cells across each sample, we performed normalization and variance stabilization using residuals from negative binomial regression with the proportion of mitochondrial genes as a covariate, as described by Choudhary *et al.* [35], using the SCTransform (SCT) version 2 function in Seurat with the glmGamPoi method [36]. A principal component analysis (PCA) dimensionality reduction was run on each sample, and the number of statistically significant principal components (PCs) was identified. Using generated artificial doublets integrated into the data set at a proportion of 0.25, the proportion of artificial nearest neighbors (pANN) was determined for each PC neighborhood size (pK). Cells with the highest pANN values were identified as predicted doublets and were removed from each sample using the doubletFinder function in the DoubletFinder package (version 2.0.4).

### Sample Integration and Clustering and Differential Gene Expression Analysis

To perform an integrative analysis of shared T cell phenotypes across each organ, samples from each mouse were combined into lists (Pancreas: 3x Isotype, 2x anti-CD226 mAb; pLN: 3x Isotype, 5x anti-CD226 mAb) to select 3150 integration features not including TCRα/β variable genes and ribosomal genes before using the SelectIntegrationFeatures function in Seurat. Using the Seurat v4 integration workflow, the SCTransform residuals for each integration feature were used to create integrated Seurat objects for each organ. Cells with related transcriptomic profiles from each organ were clustered using the first 30 PCs in the integrated assay. Briefly, the FindNeighbors function in Seurat was used to generate a nearest neighbor graph before identifying clusters by the Louvain method of community detection with a resolution of 1.0, using the FindClusters function in Seurat. Differential gene expression (DGE) between clusters was evaluated on the RNA assay using a Wilcoxon Rank Sum test. Faster implementation was achieved using the Presto package (v. 1.0.0), executed with the FindAllMarkers function in Seurat. Clusters with gene expression profiles suggestive of other immune subsets besides T cells (e.g., B lymphocytes, eosinophils), apoptotic cells, or epithelial cells were excluded from the dataset using the subset function in Seurat. Following the exclusion of cellular debris and contaminating subsets, cells were re-clustered, as described above, based on the first 13 PCs with a resolution of 0.77 for the pancreas and the first 10 PCs with a resolution of 0.7 for the pLN. This yielded 15 T cell clusters from the pancreas with 17,617 cells and 12 from the pLN with 54,229 cells. Differentially expressed genes between treatment conditions within clusters were assessed using the Wilcoxon Rank Sum test with Bonferroni correction with p_val_adj ≤ 0.05 (pLN: **Table S3**; pancreas: **Table S4**). Cell cluster annotations were assigned using *a priori* knowledge and relevant literature based on differential gene expression between clusters (**Table S5**).

### Differential Abundance Analysis

To identify differences in T cell cluster abundance between treatment conditions, we performed a differential abundance (DA) analysis for both the pancreas and pLN [37, 38]. Briefly, the edgeR package (version 3.42.4) was used to fit negative binomial generalized linear models (NB GLM). Quasi-likelihood (QL) dispersions were calculated from GLM deviances before P-values were determined using the glmQLFTest function in the edgeR package. The log fold-change (logFC) was determined based on the DA of anti-CD226 mAb-treated mice normalized to isotype-treated mice.

### TCR Repertoire Analysis & Mapping of IGRP-Reactive Clonotypes

To evaluate differences in TCR repertoires between treatment conditions, the scRepertoire package (version 2.0.0) was used to analyze V(D)J libraries. Briefly, clones were assembled by associating contigs with single-cell barcodes using the combineTCR function before merging the clonal information with the processed scRNA-seq data using the combineExpression function. To identify IGRP-reactive T cells, the TCR-11 and TCR-β CDR3 sequences were compared to the CDR3 sequences of NOD IGRP_206-124_-specific CD8^+^ T cells previously described by Kasmani *et al.* [39] using the Biostrings package (version 2.68.1). CDR3 sequences within one amino acid mismatch of either the TCR-11 or TCR-β chain of the previously published sequences were mapped as IGRP-reactive.

### Data Visualization and Statistical Analysis

Single-cell data were processed and visualized using the following R packages: Seurat [40], dsb [41], DoubletFinder [42], Presto [43], edgeR [44], SingleCellExperiment (version 1.22.0; [45]), scRepertoire [46], Biostrings [47], DittoSeq (version 1.12.2; [48]), and EnhancedVolcano (version 1.18.0; [49]). Statistical analyses were performed using GraphPad Prism software (version 9.2.0; San Diego, CA, USA) for all other data. Unless otherwise stated, flow cytometric data were analyzed by two-way ANOVA, and ELISA data were analyzed by one-way ANOVA, with Bonferroni’s post hoc test for multiple testing correction. *In vitro* suppression and chromium-release assay curves were also analyzed by two-way ANOVA with Bonferroni’s post hoc test for multiple testing correction, with AUC values compared using paired t-tests [50]. A chi-square test was used to compare insulitis severity, and a Fisher’s Exact test was used to identify differences in IGRP-reactive pancreatic CD8^+^ T cells by TCR-seq. The log-rank (Mantel-Cox) test was used to identify significant differences in disease incidence between treatment groups and calculate a hazard ratio (HR). P-values ≤ 0.05 were considered significant.

## Results

### Validation and Persistence of Anti-CD226 Blocking mAb

We validated the binding of rat 11-mouse CD226 blocking mAb (clone 480.1) by assessing the accessibility of CD226 on T cells using a fluorophore-conjugated anti-CD226 mAb (clone TX42.1) with spectral flow cytometry following *in vitro* and *in vivo* experiments. We observed significantly reduced accessibility of CD226 on CD8^+^ T cells (**Figure S7A-F**), CD4^+^ Tregs (**Figure 7G-L**), and CD4^+^ Tconv (**Figure S7M-R**) following anti-CD226 blockade relative to IgG2a isotype control, in splenocytes treated *in vitro* at a saturating concentration as well as during *ex vivo* analysis of splenic and pancreatic T cells of mice treated five weeks prior. Our *ex vivo* analysis also identified increased levels of rat IgG2a on splenic CD8^+^ T cells and NK cells of anti-CD226 treated mice (**Figure S8A-B**), suggesting the anti-CD226 blocking mAb persists for at least five weeks *in vivo*.

### CD226 Blockade Inhibits T Cell Proliferation and Promotes an Immunoregulatory Cytokine Profile

We sought to determine whether mAb blockade of CD226 signaling altered the proliferation of murine CD4^+^ or CD8^+^ T cells. We observed significant decreases in proliferation for CD4^+^ T cells blocked with anti-CD226 following either 11-CD3/11-CD28 (0.81-fold, p=0.0125) or 11-CD3/CD155-Fc (0.73-fold, p=0.048) *in vitro* stimulation relative to the IgG2a isotype control condition (**Figure 1A-B**). Similarly, CD8^+^ T cell proliferation was also significantly reduced in anti-CD226 versus isotype control treated cells following either 11-CD3/11-CD28 (0.59-fold, p=0.0015) or 11-CD3/CD155-Fc (0.40-fold, p=0.0028) stimulation (**Figure 1C-D**). To further determine whether CD226 blockade alters cytokine production, we quantified IFN-γ and IL-10 production by ELISA in the proliferation assay supernatants following 48 hours of either 11-CD3/11-CD28 or 11-CD3/CD155-Fc co-stimulation (**Figure S9**). We observed significant decreases in IFN-γ production (0.59-fold, p=0.025) and increased IL-10 production (1.20-fold, p=0.0040) following 11-CD3/CD155-Fc stimulation in anti-CD226 treated cells relative to the isotype control condition (**Figure 1E-F**).

**Figure 1.**
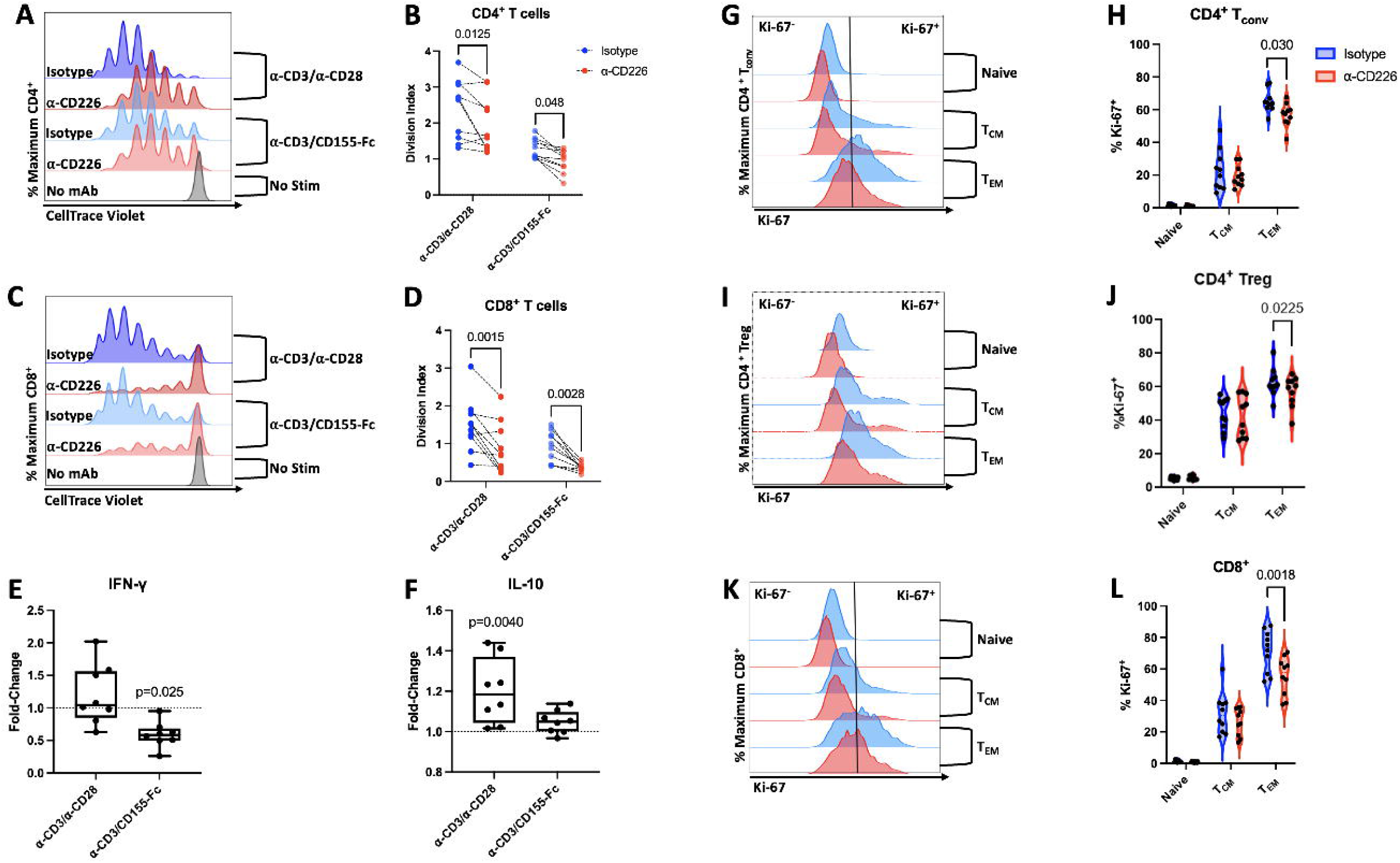
Anti-CD226 mAb Blockade Inhibits CD4^+^ and CD8^+^ T Cell Proliferation and Promotes IL-10 Production. Proliferation of (**A-B**) CD4^+^ and (**C-D**) CD8^+^ T cells within CellTrace Violet-labeled whole splenocytes was assessed by flow cytometry following *in vitro* incubation with either IgG2a isotype control (blues) or anti-CD226 mAb (reds) and subsequent stimulation. (**A, C**) Representative dye dilution plots depicting proliferation of cells treated with isotype control or anti-CD226 mAb following either no stimulation (grey) or 96 hours of plate-bound 11-CD3/11-CD28 (darker blue/red) or 11-CD3/CD155-Fc stimulation (lighter blue/red), with (**B, D**) paired dot plots showing division indices for each treatment and stimulation condition (biological n=10/condition). To assess the impact of anti-CD226 blockade on cytokine secretion, NOD splenocytes were incubated with anti-CD226 mAb or IgG2a isotype control, then stimulated for 48 hours *in vitro* with 11-CD3/11-CD28 or 11-CD3/CD155-Fc. Cell culture supernatants were analyzed by ELISA, and box and whisker plots show the fold-change in the production of (**E**) IFN-γ and (**F**) IL-10 by anti-CD226 treated relative to IgG2a treated splenocytes (biological n=8/condition). Five weeks after *in vivo* isotype control (blue) or anti-CD226 mAb (red) treatment of NOD mice, the proliferation (percent Ki67^+^) of naïve (CD44^-^ CD62L^+^), T central memory (T_CM;_ CD44^+^CD62L^+^), and T effector memory (T_EM;_ CD44^+^CD62L^-^) subsets of (**G-H**) CD4^+^ Tconv, (**I-J**) CD4^+^ Treg, and (**K-L**) CD8^+^ T cells was assessed from splenocytes by flow cytometry as shown in (**G, I, K**) representative histograms and (**H, J, L**) violin plots (biological n=10/condition). Significant P-values are reported for paired samples using (**B, D, H, J, L**) two-way ANOVA or (**E, F**) one-way ANOVA, each with Bonferroni correction for multiple comparisons.

To ascertain the impact of anti-CD226 on T cell proliferation *in vivo*, we assessed the proliferation marker, Ki-67, in isolated splenocytes five weeks post-treatment with mAb or isotype control. Interestingly, anti-CD226 blockade was associated with decreased Ki-67 expression in the T effector memory (T_EM;_ CD44^+^CD62L^-^) subsets of CD4^+^ Tconv (0.87-fold, p=0.030; **Figure 1G-H**), CD4^+^ Treg (0.92-fold, p=0.0225; **Figure 1I-J**) and CD8^+^ T cells (0.78-fold, p=0.0018; **Figure 1K-L**). Next, we assessed the effect of anti-CD226 blockade on CD4^+^ and CD8^+^ T cell frequencies across various immune and type 1 diabetes-relevant tissues five weeks after treatment (**Figure 2A**). While anti-CD226 did not significantly alter the frequencies of CD4^+^ and CD8^+^ T cells in the pancreas **(Figure 2B-C**) or thymus (**Figure 2D-E**), we identified decreased frequencies of CD8^+^ T cells in the spleen (0.79-fold, p=0.0098; **Figure 2F-G**) and pLN (0.89-fold, p=0.014, **Figure 2H-I**) of mice treated with anti-CD226. Altogether, these data indicate that inhibition of co-stimulation through CD226 promotes a shift in cytokine production toward immunoregulation and constrains the proliferation of both CD4^+^ and CD8^+^ T cells, especially in the T_EM_ compartment.

**Figure 2.**
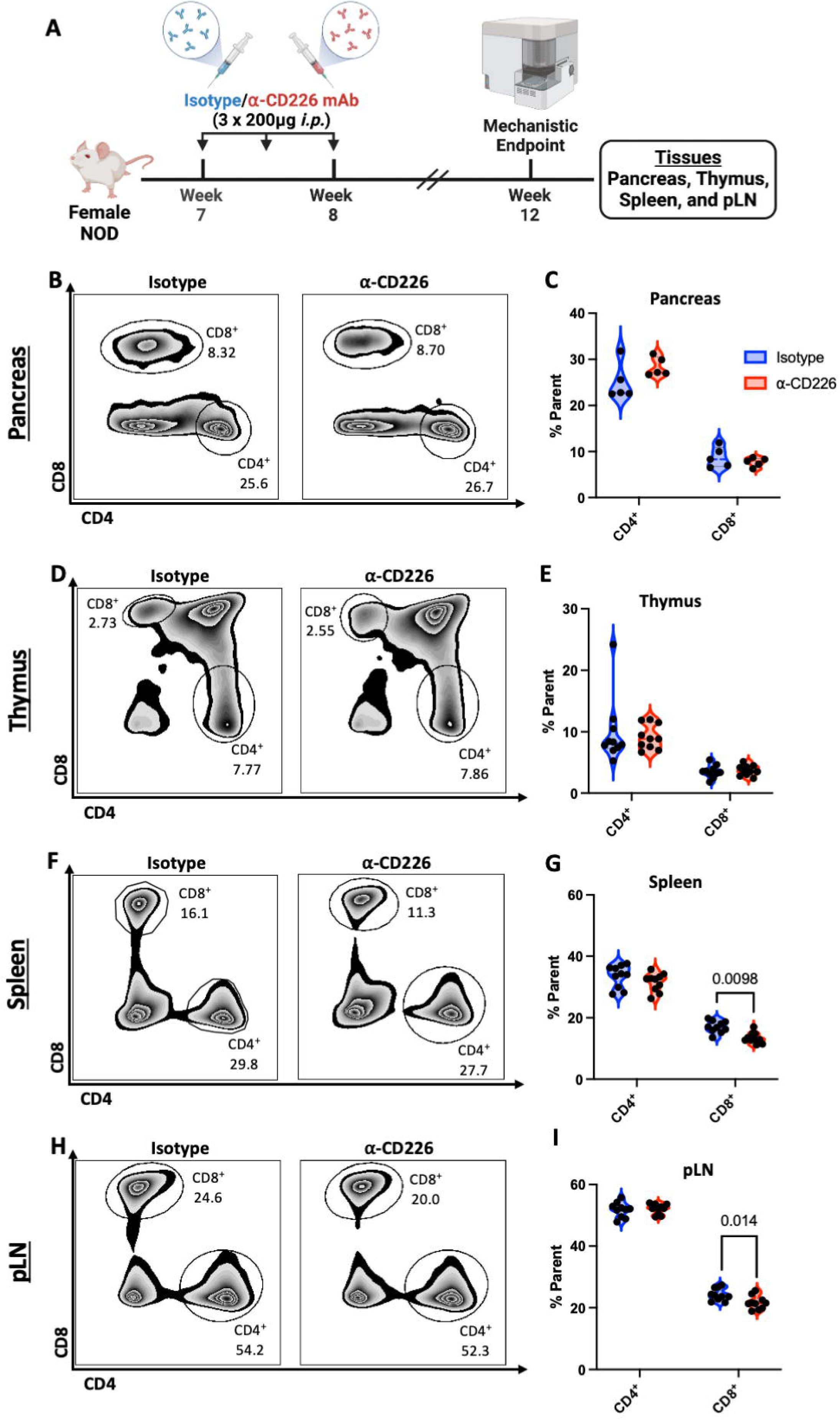
Treatment of NOD mice with anti-CD226 Reduces CD8^+^ Frequencies in Spleen and pLN. (**A**) Experimental scheme depicting the treatment of female NOD mice with anti-CD226 mAb (red) or isotype control (blue) with tissues harvested five weeks later for flow cytometric assessment (created using BioRender). Representative histograms and violin plots show the frequencies of CD4^+^ and CD8^+^ T cells in the (**B-C**) pancreas (n=5/treatment group), as well as the (**D-E**) thymus, (**F-G**) spleen, (**H-I**) and pancreatic-draining lymph nodes (pLN) (n=10/treatment group). Significant P-values are reported for two-way ANOVA with Bonferroni correction for multiple comparisons.

### CD226 Blockade Decreases Pancreatic Inflammation and Reduces Diabetes Incidence

To evaluate the *in vivo* safety and efficacy of anti-CD226 to combat the development of autoimmune diabetes, we performed an intervention study using female NOD mice treated with either IgG2a isotype control or anti-CD226 mAb between 7-8 weeks of age (**Figure 3A**). There were no significant differences between treatment groups for weight gain during the study (**Figure S10**), and no adverse complications were observed due to mAb treatment.

**Figure 3.**
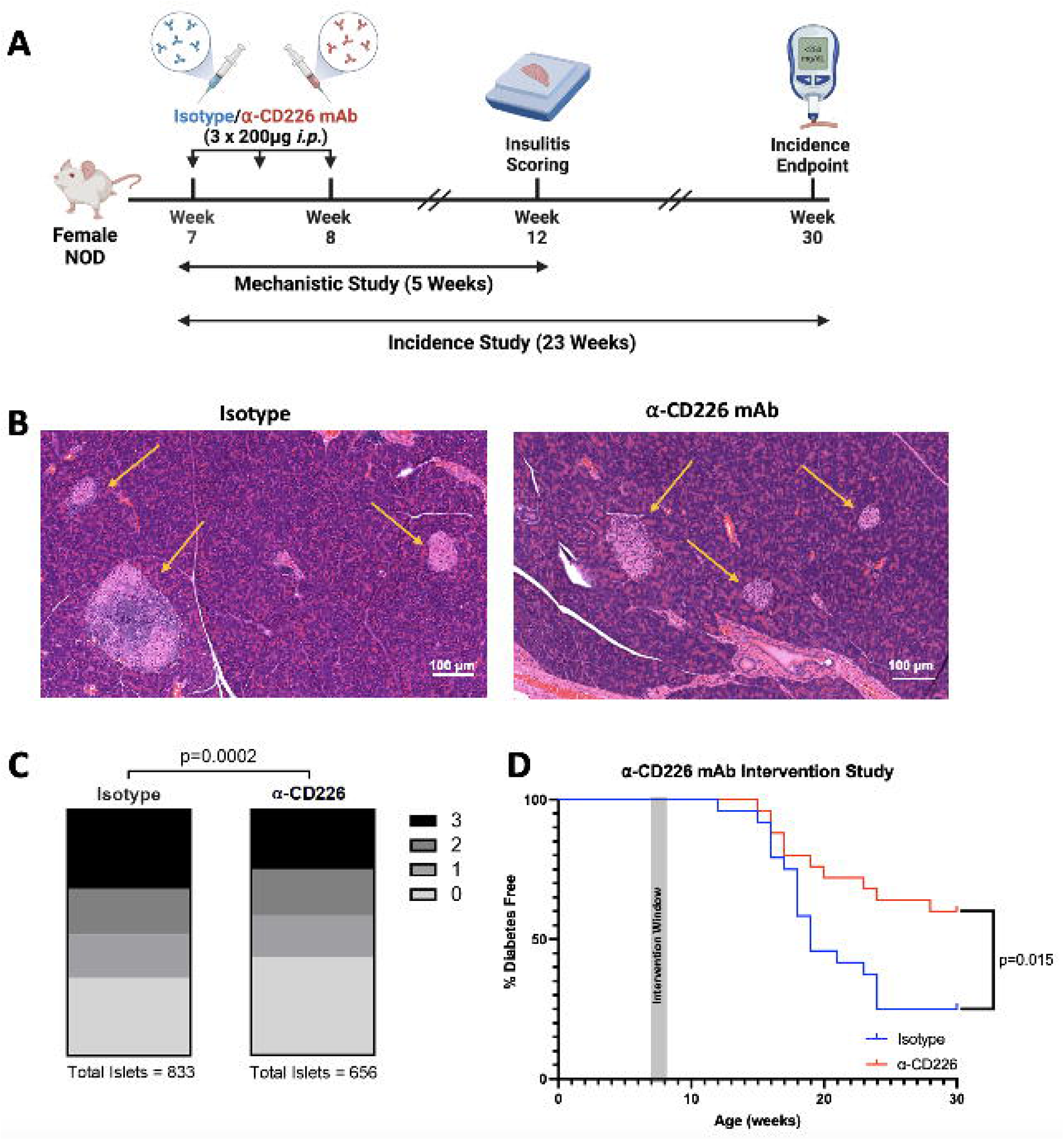
Treatment of NOD Mice With anti-CD226 mAb Reduces Spontaneous Diabetes Incidence. (**A**) Experimental scheme: insulitis severity and diabetes incidence were assessed in female NOD mice following treatment with IgG2a isotype control or anti-CD226 mAb (BioRender). (**B**) Representative images show H&E stained pancreas sections from 12-week-old mice five weeks after treatment. Islets are indicated with yellow arrows. (**C**) Stacked bar graphs show islets categorized by insulitis score (0-3) for each treatment condition (figure reports total number of islets evaluated from biological *n*=10 mice, isotype; *n*=8, anti-CD226). Significant P-value reported for Chi-Square Test. (**D**) Impact of anti-CD226 mAb (red) treatment on the incidence of autoimmune diabetes as compared to isotype control (blue) (n=25/treatment group). Significant P-value reported for Log-Rank (Mantel-Cox) test.

To understand how CD226 blockade may regulate immune infiltration in the context of autoimmune diabetes pathogenesis, we conducted a histological examination of H&E-stained pancreatic tissue sections from 12-week-old female NOD mice five weeks after administration of either IgG2a isotype control or anti-CD226 mAb (**Figure 3A-B**). We observed a reduced severity of insulitis in the anti-CD226 treated group (0.84-fold, p=0.0002, **Figure 3C**). Crucially, we observed that NOD mice treated with anti-CD226 had a significantly reduced disease incidence (Hazard Ratio=0.41, p=0.015) relative to mice treated with the isotype control (**Figure 3D**).

### CD226 Blockade Augments Treg Suppressive Capacity

We next sought to understand the mechanism of protection afforded by anti-CD226 treatment. Our previous work in NOD mice and primary human T cells showed that CD226 signaling antagonized Treg stability and functionality [8, 23, 51], which are necessary for regulating immune tolerance. One aspect critical to the function of Tregs is adequate IL-2 receptor (IL-2R) signaling, with previous work by Permanyer *et al*. demonstrating that a partial loss of IL-2R signaling impairs the Treg lineage stability and suppressive function required to inhibit autoimmunity [52]. Therefore, we assessed whether CD226 blockade impacted Tregs *in vivo*, using flow cytometry to examine the expression of the high-affinity IL-2R 11-chain (CD25) on Foxp3^+^Helios^+^ thymic Tregs (tTregs) across a variety of NOD tissues (**Figure 4A-H**). We identified significantly higher expression of CD25 on tTregs of anti-CD226 treated mice in all organs examined, including the thymus (1.12-fold, p=0.0014; **Figure 4A, 4E**), spleen (1.10-fold, p=0.015; **Figure 4B, 4F**), pLN (1.05-fold, p=0.015; **Figure 4C, 4G**), and pancreas (2.05-fold, p=0.0073; **Figure 4D, 4H**) as compared to isotype-treated mice. To validate this finding, we measured IL-2-induced STAT5 phosphorylation in Tregs using splenocytes from mice administered anti-CD226 or IgG2A five weeks prior. We observed that Tregs from anti-CD226 treated animals demonstrated greater phosphorylation of STAT5 after 60 minutes of co-culture with rhIL-2 (1.39-fold, p=0.0007; **Figure 4I-L**). To understand how CD226 blockade may impact Treg functionality, we performed *in vitro* suppression assays and observed that NOD Tregs pre-treated *in vitro* with anti-CD226 demonstrated significantly increased suppression of CD4^+^ Tconv responders compared to Tregs pre-treated with the isotype control (**Figure 4M-O**). These findings suggest that anti-CD226 may augment Treg activation and suppressive capacity.

**Figure 4.**
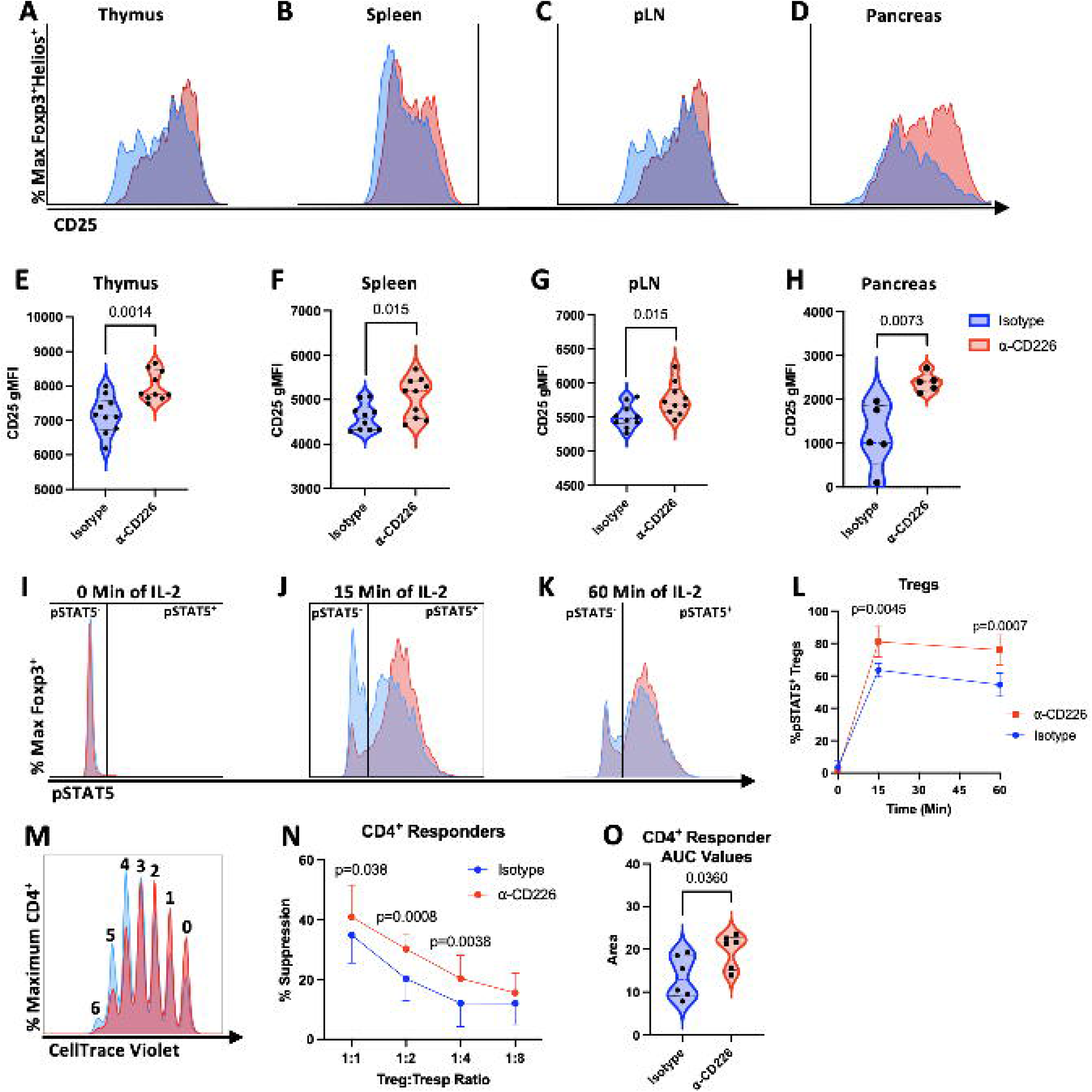
Anti-CD226 mAb Blockade Augments Treg Suppressive Capacity. CD25 expression was measured by flow cytometry in various tissues from female NOD mice five weeks after treatment with either IgG2a isotype control (blue) or 11-CD226 mAb (red). (**A-D**) Representative histograms and (**E-H**) violin plots show the geometric mean fluorescence intensity (gMFI) of CD25 on Foxp3^+^Helios^+^ tTregs in the (**A, E**) thymus, (**B, F**) spleen, and (**C, G**) pancreatic-draining lymph nodes (pLN) (n=10/treatment group), as well as (**D, H**) pancreas (n=5/treatment group). Significant P-values are reported for unpaired t-tests. (**I-L**) Phosphorylation of STAT5 (pSTAT5) following *in vitro* IL-2 stimulation was measured by flow cytometry in whole splenocytes obtained from 12-week-old female NOD mice, five weeks after treatment with either IgG2a isotype control (blue) or 11-CD226 mAb (red). Representative histograms show staining for pSTAT5 in CD4^+^Foxp3^+^ Tregs following (**I**) 0, (**J**) 15, or (**K**) 60 minutes of stimulation of IL-2 with (**L**) pSTAT5^+^ Treg frequencies over 60 minutes of IL-2 stimulation (n=4/condition). Significant P-values are reported for two-way ANOVA with Bonferroni correction for multiple comparisons. (**M**) Representative dye dilution plot shows the proliferation of CTV labeled CD4^+^ T responders (Tresp) in an *in vitro* suppression assay at the 1:2 Treg:Tresp ratio following Treg incubation with IgG2a isotype control (blue) or 11-CD226 mAb (red). (**N**) Percent suppression of CD4^+^ Tresp was determined using the division index method at each Treg:Tresp ratio (n=6/condition, significant P-values reported for two-way ANOVA with Bonferroni correction), with additional comparisons made using (**O**) area under the curve (AUC) values for each suppression curve (paired t-test).

### CD226 Blockade Reduces Effector T Cell Cytotoxicity

The autoimmune destruction of pancreatic beta-cells by cytotoxic CD8^+^ T cells is a hallmark of type 1 diabetes pathogenesis and an important target for therapeutic interventions [53]. Our prior work in *Cd226* KO NOD mice demonstrated that diabetes protection was associated with reduced TCR affinity of pancreatic CD8^+^ T cells specific for the type 1 diabetes autoantigen, IGRP [20]. Thus, we quantified the frequency of IGRP reactive CD8^+^ T cells in the pancreata of 12-week-old female NOD mice five weeks after administration of either IgG2a isotype control or anti-CD226 mAb. Using IGRP_206-214_ tetramer staining (**Figure 5A-B**), we identified that NOD mice treated with anti-CD226 had a significantly reduced frequency of IGRP-reactive CD8^+^ T cells (0.50-fold, p=0.0317) infiltrating the pancreas. To further determine whether anti-CD226 mAb can directly influence the cytotoxic activity of CD8^+^ T cells to protect pancreatic beta-cells, we utilized an *in vitro* model of cell-mediated lysis (CML) (**Figure 5C**) leveraging the murine AI4 (Vα8/Vβ2)-autoreactive CD8^+^ T cell clone, which recognizes a peptide derived from the insulin A chain (InsA_14-20_) in the context of H-2D^b^ [32]. We observed that treatment of AI4-CD8^+^ T cells with anti-CD226 before antigen-specific activation reduced specific lysis of the murine NIT-1 pancreatic beta-cell line compared to NIT-1 cells co-cultured with isotype control-treated AI4-CD8^+^ T cells (**Figure 5D-E**). These data suggest that anti-CD226 may limit autoreactive effector T cell frequency and cytotoxicity within the target organ, collectively contributing toward reduced beta-cell destruction.

**Figure 5.**
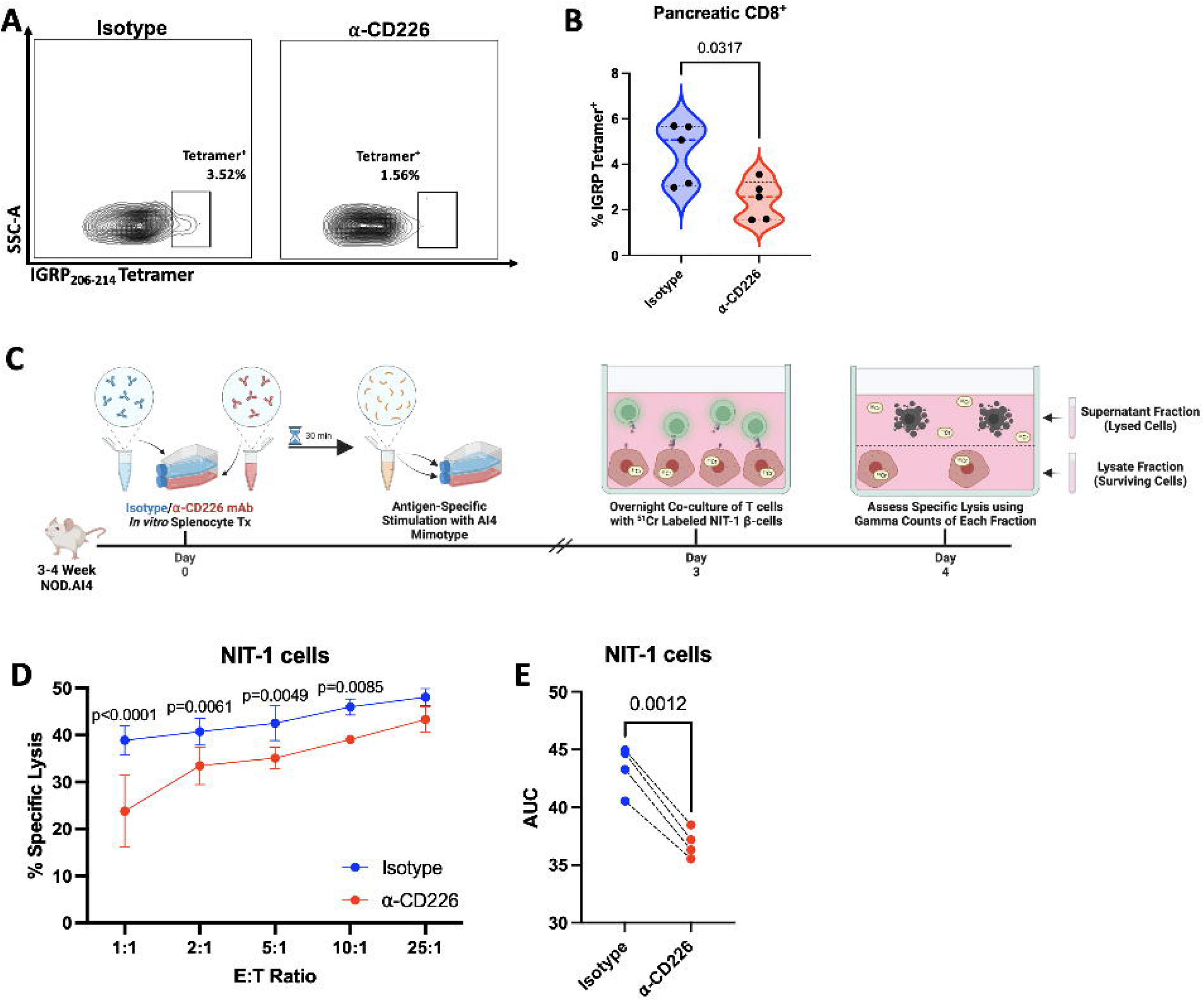
Anti-CD226 mAb Blockade Diminishes T Cell Cytotoxicity. CD8^+^ IGRP-reactive T cells were quantified using IGRP_206-214_ tetramer staining in lymphocytes isolated from the pancreases of 12-week-old female NOD mice five weeks after treatment with isotype control (blue) or anti-CD226 (red). (**A**) Representative flow cytometric contour plots and (**B**) violin plots show the frequency of IGRP tetramer^+^ CD8^+^ T cells (n=5/treatment group). Significant P-value reported for unpaired t-test. (**C**) Experimental scheme depicting CML assay to assess the effect of CD226 blockade on the cytotoxic activity of autoreactive AI4-mouse lymphocytes killing NIT-1 pancreatic beta-cells, as measured by ^51^Cr-release (created with BioRender). (**D**) Percent specific lysis at each Effector:Target cell (E:T) ratio (n=4/condition, significant P-values reported for two-way ANOVA with Bonferroni correction for multiple comparisons) with (**E**) AUC values (paired t-test).

### CD226 Blockade Modulates Expression of Genes Associated with T cell Activation

To determine the transcriptional networks underlying the above-described phenotypic and functional changes induced by anti-CD226 blockade, we used scRNA-seq to characterize gene expression profiles in T cells isolated from the pLN and pancreas of anti-CD226 and isotype control-treated NOD mice (**Figure 6A**). Within the 12 pLN T cell clusters identified (**Figure 6B**), we observed increased expression of genes associated with migration, chemotaxis, and tissue retention (**Figure 6C** and **Table S3**), including *Cxcr4* (CD4^+^ naïve, CD8^+^ naïve, CD8^+^ T resident memory (T_RM_), CD8^+^ cycling), *Il16* (CD8^+^ naïve, T_CM_, CD8^+^ cycling), and *Cd69* (CD4^+^ naïve, CD4^+^ T_EM_, CD8^+^ naïve, CD8^+^ T_CM_) among mice treated with anti-CD226, compared to the isotype control. We also observed increased expression of cytokine receptors, *Il7r* (CD8^+^ naïve, T_CM_, T_RM_, cycling and CD4^+^ naïve, T_CM_, T_EM_) and *Ifngr1* (CD8^+^ T_CM_, T_RM_, cycling and CD4^+^ T_CM_), as well as the transcription factor *Stat1* (CD8^+^ naïve, T_CM_, T_RM_, cycling and CD4^+^ naïve, T_CM_, T_EM_, resting) in anti-CD226 treated mice. None of these genes were differentially expressed in Treg, γδ T cell, or activated CD8^+^ T cell clusters. Across most of the 15 pancreas T cell clusters identified (**Figure 6D**), we observed increased transcription of *Cdk8*, encoding for cyclin-dependent kinase 8 and *Il31ra*, which encodes IL-31Ra, known to be involved in Th2 responses, as well as decreases in *Dock2*, which is associated with both T cell activation and trafficking through regulating the actin cytoskeleton (**Figure 6E** and **Table S4**), in mice treated with anti-CD226. The anti-CD226 group also exhibited reduced expression of *Il7r* (CD4^+^ naïve, cycling), *Txnip* (CD4^+^ cycling), *Cd69* (CD8^+^ T_EM_), and *Ifngr1* (Th2). Additionally, we observed increases in *Smad7* (CD4^+^ T_EM_, T_CM_, tTreg and CD8^+^ naïve) following anti-CD226 treatment. Together, these data indicate that anti-CD226 treatment decreased the expression of genes associated with activation and increased the expression of genes related to migration. While we did not observe any significant differences in T cell cluster abundance in the pancreas (**Figure S11** and **Table S5**), we saw a reduced abundance of the CD8^+^ T_CM_ and CD8^+^ cycling clusters in the pLN (**Figure S12** and **Table S5**) among mice treated with anti-CD226, suggesting the reduced frequency of circulating CD8^+^ T cells observed by flow (**Figure 2**) may be associated with reductions in activated T cells.

**Figure 6.**
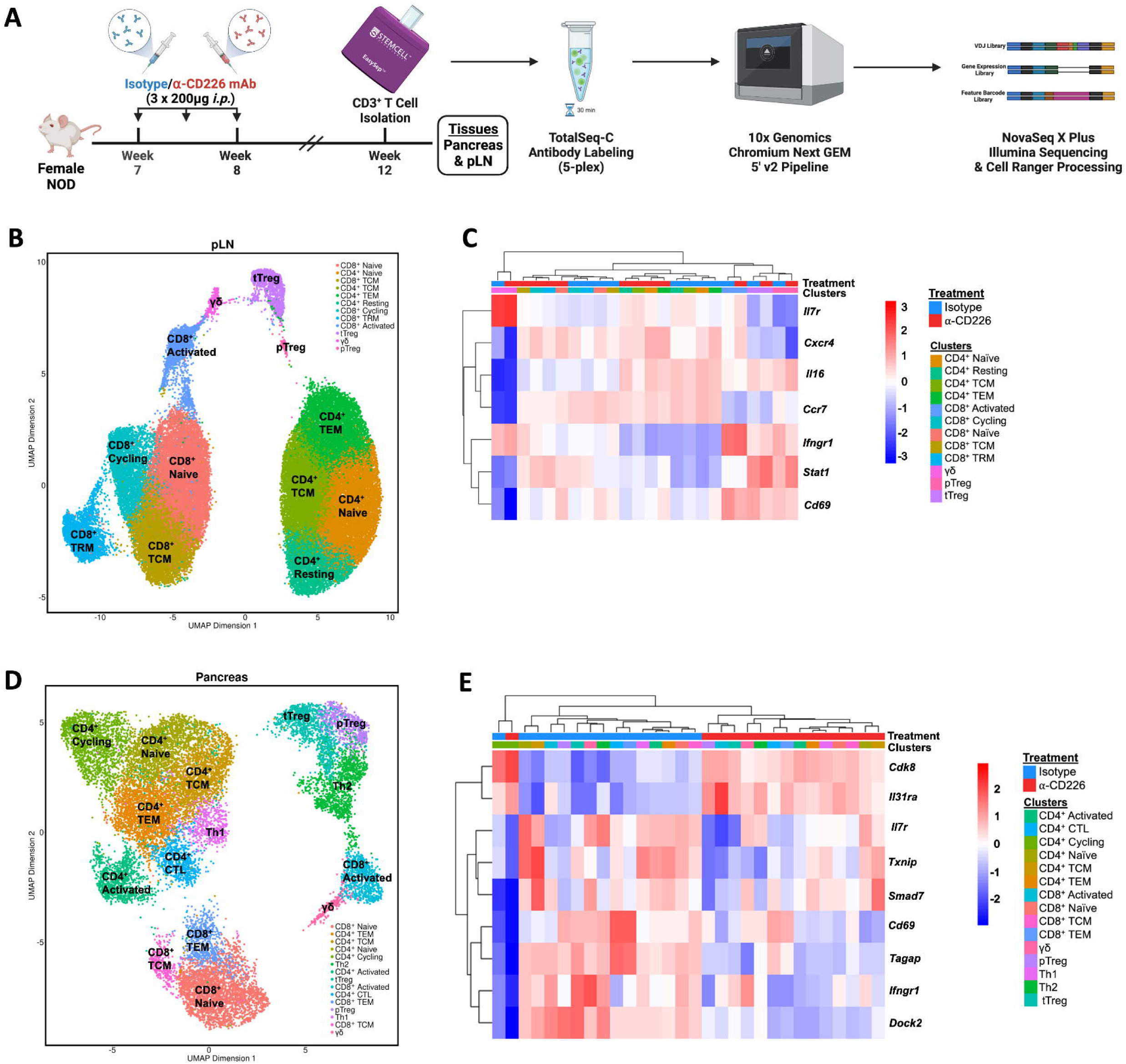
Anti-CD226 mAb Blockade Reduces Expression of T cell Activation Genes. To identify differentially expressed genes as a result of anti-CD226 blockade, (**A**) scRNA-seq/CITE-seq was performed on CD3^+^ T cells isolated from the (**B-C**) pLN (n=3 isotype, n=5 anti-CD226 mAb) and (**D-E**) pancreas (n=3 isotype, n=2 anti-CD226) of 12-week-old female NOD mice, five weeks after anti-CD226 or isotype administration. (**A**) Experimental scheme (created with BioRender). (**B, D**) Uniform Manifold Approximation and Projection (UMAP) plots annotated with distinct T cell clustering for each organ assessed (pLN = 54,229 cells; pancreas = 17,617 cells). (**C, E**) Heatmaps show aggregated gene expression Z-scores (pLN: *Il7r, Cxcr4, Il16, Ccr7, Ifngr1, Stat1, Cd69*; pancreas: *Cdk8, Il31ra, Il7r, Txnip, Smad7, Cd69, Tagap, Ifngr1, Dock2*) by cluster and treatment condition.

### CD226 Blockade Regulates T Cell Clonal Expansion

Co-stimulatory signaling through CD226 enhances T effector function and cytotoxicity [54], and in settings of cancer, CD226 signaling has been leveraged to promote clonal expansion of anti-tumor CD8^+^ T cells [55]. To evaluate how anti-CD226 blockade might regulate autoreactive T cell clonal expansion in autoimmune diabetes, we performed single-cell TCR sequencing (scTCR-seq) to annotate and quantify TCR clonality in the pancreas and pLN of anti-CD226 versus isotype-treated NOD mice. The most extensive degree of clonal expansion in the pancreas was localized in an activated CD8^+^ cluster **(Figure 7A**), with those expanded activated CD8^+^ T cell clones demonstrating reduced expression of the pro-inflammatory associated genes, *Gzma* (log_2_FC: -53.7, FDR-adjusted p=8.65e^-11^; [56]) and *Xcl1* (log_2_FC: -82.7, FDR-adjusted p=5.45e^-5^; [57]) in anti-CD226 mAb relative to isotype treated mice (**Figure 7B**). Although we observed greater amounts of “medium” (0.001 < X < 0.01) and reduced levels of “small” (1e^-4^ < X < 0.001) expanded clone cutoffs following anti-CD226 treatment (**Figure 7C**), we did not observe any significant differences in the overall abundance of small and medium expanded clones by cell cluster (**Figure 7D-F**). Using previously reported CDR3 sequences [39], we found that most IGRP_206-218_-reactive T cells were localized to the activated CD8^+^ T cell cluster (**Figure 7G**). Notably, there was a significantly reduced frequency of IGRP-reactive T cells in the activated CD8^+^ T cell cluster in the pancreas from the anti-CD226 treatment group compared to isotype-treated mice (0.61-fold, p=0.022, **Figure 7H**). Upon examining the impacts of anti-CD226 on clonal expansion in the pLN, we did not observe any significant differences compared to isotype-treated mice (**Figure S13**), suggesting that the effects of anti-CD226 on T cell priming in the pLN may be more apparent at the site of inflammation in the pancreas, where autoreactive T cells are undergoing proliferation [58].

**Figure 7.**
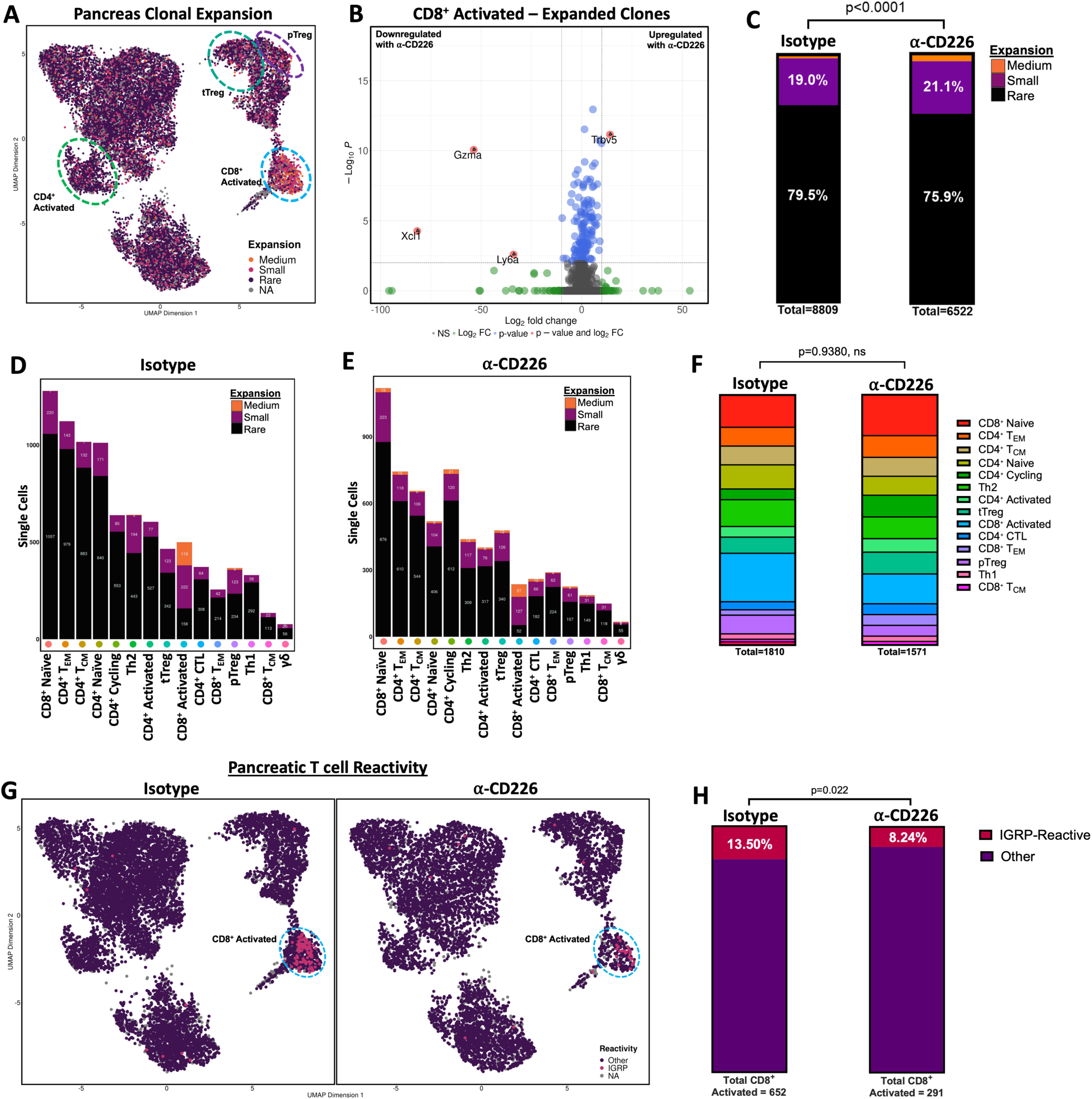
Anti-CD226 mAb Blockade Alters T Cell Clonal Expansion in the Pancreas. To determine the impact of anti-CD226 blockade on T cell clonal expansion, TCR-seq data was merged with corresponding scRNA-seq data on T cells isolated from 12-week-old female NOD pancreas five weeks after treatment with IgG2a isotype control or anti-CD226 mAb. (**A**) The degree of clonal expansion within each cluster was assessed by overlaying the proportion of each clonotype (Medium: 0.001 < X < 0.01 (orange), Small: 1e^-4^ < X < 0.001 (pink), Rare: 0 < X < 1e^-4^ (purple), NA: no CDR3 sequence for cell barcode (gray)) onto a UMAP projection of T cells. (**B**) Volcano plot shows differentially expressed genes of small and medium expanded clones in the highly expanded cluster of activated CD8^+^ T cells. Genes with Log_2_ fold change (FC) < 10 and Log_10_ P-value > 1.3 are shown in gray (not significant, NS). Genes with Log_2_ FC > 10 but Log_10_ P-value > 1.3 are shown in green. Genes with Log_2_ FC < 10 but Log_10_ P-value < 1.3 are shown in blue. Six genes were identified as having significant differential expression (Log_10_ P-value < 1.3 and Log_2_ FC > 10) shown in red with annotations. Stacked bar plots show that (**C**) while there is an increased frequency of medium-expanded clones following α-CD226, the distributions of clonal expansion by cluster in mice treated with (**D**) isotype (**E**) or anti-CD226 remain (**F**) unchanged when comparing the overall distribution of small and medium clones by cluster. (**G**) UMAP plots split by treatment condition (isotype = 10,077 cells; anti-CD226 = 7,540 cells) show T cells containing CDR3 sequences within one amino acid of previously published IGRP-reactive clones, annotated as IGRP-reactive (pink) or having any other reactivity (purple). (**H**) Stacked bar plots show the frequency of IGRP-reactive clones within the activated CD8^+^ T cell cluster between isotype (n=3) or anti-CD226 treated mice (n=2) with the total number of activated CD8^+^ T cells per treatment group shown below. Significant P-value reported for Fisher’s Exact Test.

## Discussion

Treatment of pre-diabetic NOD mice with an anti-CD226 blocking mAb resulted in significant prevention of autoimmune diabetes and insulitis, representing an important advance toward the translation of new drug candidates targeting this pathway for the treatment of human type 1 diabetes. This builds on prior knowledge of *CD226* as a candidate risk gene [59], as well as our prior work using global and Treg-specific KO NOD mouse strains where we have previously reported that CD226 plays a significant role in the development of spontaneous autoimmune diabetes by controlling peripheral T cell activation [20] as well as Treg function and stability [23]. Furthermore, our human studies have revealed that CD226^+^ Tregs demonstrate increased IFN-γ production and decreased lineage stability [8]. In contrast, isolation and *ex vivo* expansion of CD226^−^ Tregs yields a more lineage-stable and suppressive Treg population [51]. Given these findings and to advance the translation of new drug candidates for the prevention and treatment of human type 1 diabetes, we investigated whether mAb-mediated blockade of CD226 could reduce diabetes incidence in NOD mice by reducing effector T cell co-stimulatory signaling and improving Treg functionality.

CD226 co-stimulation is known to contribute to both T cell activation and proliferation [17], and we confirmed that *in vitro* blockade with anti-CD226 mAb reduced both 11-CD3/11-CD28 and 11-CD3/CD155-Fc stimulated T cell proliferation. Moreover, upon *ex vivo* analysis of splenocytes from anti-CD226-treated NOD mice, we identified reduced proliferative capacity of both CD4^+^ and CD8^+^ T cells in the T_EM_ compartment compared to the isotype control-treated group. Our previous work has shown that human CD4^+^ T_EM_ have the highest expression of CD226 compared to other CD4^+^ T cell subsets [8]; therefore, a potential explanation for this finding is that anti-CD226 may preferentially suppress T_EM_ compared to naïve T cells.

Considering that the expansion of insulin-specific CD4^+^ T_EM_ has been correlated with insulin autoantibody-positivity in recent-onset stage 3 type 1 diabetes [60], along with the known propensity of CD8^+^ T_EM_ to have substantial cytotoxic function [61], our findings support pharmacotherapeutic targeting of CD226 signaling as a means to constrain activation of diabetogenic effector T cell subsets.

Interestingly, while anti-CD226 clone 480.1 is only thought to participate in blocking CD226 interactions [62], we did observe slight reductions in CD8^+^ T cell frequencies in the spleen and pLN. Previous work by our lab has shown that *Cd226* KO NOD mice have impaired thymocyte development [20]. However, we observed no differences in CD4^+^ nor CD8^+^ thymocyte frequencies as a result of anti-CD226 mAb treatment, which may be due to the timing of treatment in relation to thymic development. Hence, the reduced CD8^+^ T cell frequencies observed likely reflect the noted impacts on T cell proliferation, resulting from impeded co-stimulatory signaling (signal 2) and downstream T cell activation. Indeed, anti-CD226 appears to limit T cell activation during pro-inflammatory conditions, as evidenced by a reduction in pancreatic T cell transcription of genes associated with activation and lymphocyte trafficking, such as *Cd69* and *Dock2* [63–65]. Importantly, we observed reduced expression of *Dock2* in the activated cluster that contains putative IGRP-reactive CD8^+^ T cells. While we did not observe any significant differences in the abundance of pancreatic Th1 and Th2 clusters at this particular time point, we note a trend towards reduced pancreatic Th1 abundance and, interestingly, increased expression of the Th2-associated gene, *Il31ra,* in anti-CD226 treated mice, suggesting that CD226 blockade may be altering T cell polarization [66].

Beyond serving as a marker of activation, CD69 expression has also been associated with increased CD8^+^ T cell tissue residency [67, 68], where we observed increased expression of *Cxcr4* and *Cd69* in naïve CD4^+^, naïve CD8^+^, and CD8^+^ T_CM_ clusters from the pLN of anti-CD226-treated mice, suggesting anti-CD226 mAb may be inducing retention of T cells in secondary lymphatics. While we did not observe any differences in T cell cluster abundance in the pancreas, we noted significant reductions in the abundance of CD8^+^ cycling and CD8^+^ T_CM_ cells in the pLN of anti-CD226 versus isotype-treated mice. CD226 is involved in platelet and monocyte adhesion to endothelial cells [69, 70]; therefore, anti-CD226 may contribute to changes in T cell migratory capacity by limiting adhesion.

Previous work by Johnston et al. has demonstrated that the blockade of CD226 in a BALB/c model of colorectal carcinoma reduced IFN-γ-producing CD8^+^ frequencies and subsequently accelerated tumor growth [71]. In autoimmunity, CD226 blockade and *Cd226* KO in a C57BL/6 model of EAE by Zhang et al. showed increased production of IL-10 and diminished production of both IFN-γ and IL-17, leading to improved EAE outcomes [22]. We similarly observed an association between reduced CD226 activity and a more immunoregulatory phenotype, with anti-CD226 mAb treated cells demonstrating decreased IFN-γ and increased IL-10 production *in vitro.* Notably, CD226 blockade may be altering the expression of genes influenced by the IFN pathway *in vivo*, as evidenced by increased pancreatic T cell expression of *Cdk8*, which encodes cyclin-dependent kinase 8, in the anti-CD226 mAb treatment group. Cdk8 is reported to phosphorylate the Ser^727^ residue of STAT1 and modulate IFN-γ responsive gene expression [72]. Furthermore, we observed increased *Ifngr1* transcription in pLN T cells yet decreased *Ifngr1* transcription in pancreatic T cells of anti-CD226 treated mice, suggesting that at this particular time point, anti-CD226 mAb is inducing a more immunoregulatory phenotype in the target organ, perhaps, by limiting the migration of IFNGR1^+^ T cells.

Compared to healthy controls, HLA-A*02-01 individuals with type 1 diabetes have been reported to possess expanded populations of circulating IGRP-reactive memory CD8^+^ T cells with a distinct *TRA* motif [73]. IGRP-reactive CD8^+^ T cells have previously been shown to be involved in pancreatic beta-cell destruction and inflammatory cytokine production [74]. Therefore, diminishing cell-mediated destruction of pancreatic beta-cells by autoreactive CD8^+^ T cells is a key objective for immunotherapies seeking to restore immune tolerance in type 1 diabetes [53, 75]. We have previously reported that *Cd226* KO reduces IGRP-reactive pancreatic CD8^+^ T cells in NOD mice [20]. Importantly, herein we observed decreased IGRP-reactive CD8^+^ T cells with anti-CD226 treatment by flow-based MHC tetramer staining and scTCR-seq, as well as reductions in CML killing when insulin-specific CD8^+^ T cells were treated with anti-CD226 mAb before co-culture with NIT-1 target beta-cells. These findings support that anti-CD226 treatment may reduce autoreactivity by limiting co-stimulatory signaling and, subsequently, the clonal expansion and cytotoxicity of autoreactive T cells.

Beyond limiting the cytotoxic and pro-inflammatory potential of effector T cells, we also identified that anti-CD226 treatment promoted immunoregulatory changes in the Treg compartment. Importantly, we observed greater expression of *Smad7* in the tTreg cluster in the pLN of anti-CD226 treated mice, where the upregulation of *Smad7* has been implicated in the inhibition of PD-L2/PD-1 [76] and TGF-β [77] signaling and associated with chronic infection and autoimmunity [78]. However, long-term increases in Treg and decreases in Th17 frequencies, alongside upregulation of *Smad7*, in an Echinococcus multilocularis (EM) model of chronic infection [79] have also been observed, suggesting that anti-CD226 may be modulating Treg activity. Moreover, phosphorylation of the regulatory transcription factor, FOXO1, is thought to be mediated by PI3K signaling downstream of CD226 [80]. Ouyang et al. have shown that Foxo1^-/-^Foxo3^-/-^ mice have impaired FoxP3^+^ Treg development and function [81]. Therefore, we hypothesized that the blockade of CD226 signaling would augment Treg functionality.

Accordingly, to examine whether anti-CD226 may regulate Treg stability and functionality *in vivo*, we performed *ex vivo* phenotyping of Tregs in the spleen, thymus, pLN, and pancreas from anti-CD226 and isotype control-treated Foxp3-GFP Treg reporter NOD mice. While we did not observe significant differences in Treg frequency, anti-CD226 treated mice exhibited increased CD25 expression on Tregs in all organs examined, suggesting that anti-CD226 may increase Treg fitness by bolstering IL2R avidity for IL-2. This was further supported by phospho-flow cytometry indicating that Tregs isolated from mice treated with anti-CD226 have increased pSTAT5-signaling following stimulation with IL-2 *ex vivo*. Likewise, we have previously observed that CD226^−^ human Tregs demonstrate greater CD25 expression following *ex vivo* expansion [51], potentially indicating that less CD226 signaling results in CD25 upregulation or persistence. Previous studies have identified that Tregs exhibiting higher IL-2 avidity demonstrate improved lineage stability and function [82, 83]. Thus, we interrogated whether anti-CD226 blockade could improve Treg suppressive capacity.

Using *in vitro* suppression assays, we observed that pre-treatment of Tregs with anti-CD226 improved their suppressive capacity by reducing CD4^+^ Tresp proliferation after four days of co-culture. This finding is significant as the impairment of Treg suppressive function allows for the proliferation of autoreactive T cells and, subsequently, the pathogenesis of autoimmune diseases such as type 1 diabetes [84, 85], where these autoreactive T cells contribute to cell-mediated killing of pancreatic beta-cells. Overall, the reductions in CD8^+^ cytotoxicity and improvements in Treg suppressive capacity conferred by anti-CD226 treatment suggest that CD226 blockade may enable a shift away from cytotoxicity and towards immunoregulation in the T cell compartment.

Antibody-based immunotherapeutics represent a popular approach for treating many autoimmune diseases [86]. In the context of type 1 diabetes, a low dose of polyclonal anti-thymocyte globulin (ATG) may act by depleting T cells and inducing exhaustion, and has been shown to preserve beta-cell function in new-onset patients [87–89]. Furthermore, monoclonal antibodies can also inhibit the activation of specific immune cell subsets. For example, rituximab (11-CD20) is used to treat RA and systematic lupus erythematosus (SLE) by depleting B cells [6, 90], whereas teplizumab (11-CD3) is thought to suppress autoreactivity in type 1 diabetes by inducing exhaustion in T cells and augmenting Treg activation [91, 92]. Beta-cell replacement therapies have sought to restore the endogenous control of blood glucose; however, combining beta-cell replacement with immunotherapy may improve engraftment and promote beta-cell longevity by delaying antigen-specific killing mediated by T cells [53]. Given that activated T cells express CD226 at greater levels, we propose that anti-CD226 mAb may preferentially suppress the activity of autoreactive cells to preserve endogenous or transferred beta-cells.

Beyond the important immunoregulatory shifts observed following treatment, anti-CD226 mAb was well-tolerated in mice with no observed adverse effects. Nevertheless, while CD226 is known to be most highly expressed on effector T cells, CD226 is also expressed on many other cell subsets, including NK cells, innate lymphoid subsets, monocyte/macrophages, dendritic cells, IL-10-producing Tr1-like T cells, hematopoietic progenitor cells, endothelial cells, and platelets. Thus, anti-CD226 mAb would need to be carefully titrated to avoid disrupting critical processes such as the clearance of pathogens and tumors as well as blood clotting [17, 69, 70, 93–96]. In addition to safety studies, further investigations should determine the potential impact of CD226 genetic variants on treatment response to anti-CD226 mAb blockade as a consideration for precision medicine approaches.

Our study supports the therapeutic potential of anti-CD226 mAb to reduce spontaneous diabetes incidence and insulitis in the NOD mouse. Although it is possible that disease prevention in the NOD mouse model may not directly translate to clinical efficacy in human type 1 diabetes [97], our data demonstrate that anti-CD226 has an immunoregulatory effect on T cells by bolstering Treg and limiting effector function. Altogether, our results support the continued investigation of the CD226:TIGIT signaling axis in immune tolerance. Indeed, studies investigating TIGIT-Immunoglobulin fusion proteins as an additional approach to modulate CD226:TIGIT signaling and restore immune tolerance are ongoing in our laboratory. In summary, these findings present a novel strategy using anti-CD226 mAb as a deliverable biologic to augment Treg function, diminish effector T cell cytotoxicity, and reduce spontaneous diabetes incidence in the NOD mouse model.

## Abbreviations

APCs: Antigen-presenting cells
ATG: Anti-thymocyte globulin
CITE-Seq: Cellular indexing of transcriptomes and epitopes
CML: Cell-mediated lysis
CPM: Counts per minute
CTV: CellTrace™ Violet
DGE: Differential gene expression
DI: Division index
EAE: Experimental allergic encephalomyelitis
FC: Fold change
FMO: Fluorescence-minus one
gMFI: Geometric mean fluorescence intensity
HPCs: Hematopoietic progenitor cells
IGRP: Islet-specific glucose-6-phosphatase catalytic subunit-related protein
InsA: Insulin A chain
iTreg: Induced regulatory T cell
KO: Knockout
logFC: Log fold-change
mAbs: Monoclonal antibodies
MS: Multiple sclerosis
NBGLM: Negative binomial generalized linear models
pANN: Proportion of artificial nearest neighbors
pLN: Pancreatic-draining lymph nodes
QL: Quasi-likelihood
RA: Rheumatoid arthritis
scRNA-seq: Single-cell RNA sequencing
scTCR-seq: Single-cell T cell receptor sequencing
SLE: Systematic lupus erythematosus
Tc1: Type 1 CD8+
TCM: T central memory
Tconv: Conventional CD4+ T cell
TEM: T effector memory
Th1: T helper type 1
Treg: Regulatory T cell
Tresp: Responder T cell
TRM: T resident memory
tTregs: Thymic Tregs
UMAP: Uniform Manifold Approximation and Projection
UMI: Unique molecular identifier

## Acknowledgments

We thank Thinzar Myint and members of the Brusko Laboratory at the University of Florida Diabetes Institute for discussions and technical assistance. We also appreciate the IGRP_204-214_-BUV395 tetramers, kindly provided by Brian Fife and Justin Spanier (University of Minnesota). We thank Brandy Roldan-Martinez and Heather Bonanno (University of Florida) for mouse breeding and husbandry. We also thank the UF Molecular Pathology Core for pancreata processing and the UF Center for Immunology and Transplantation (CIT) for spectral flow cytometer and cell sorter access.

## Data Availability Statement

The single-cell sequencing data generated for this study can be found in the GEO repository, with the accession number GSE254403 or upon request.

## Author Contributions

**MEB**: Conceptualization, Methodology, Investigation, Writing – Original Draft, Visualization, Formal Analysis, Software, Data Curation, Supervision, Project Administration. **PT**: Conceptualization, Methodology, Investigation, Writing – Review & Editing, Visualization, Funding Acquisition, Supervision, Project Administration. **LDP**: Investigation, Software, Writing – Review & Editing, Funding Acquisition. **EJK, SV**, **LKS**: Investigation, Writing – Review & Editing. **ALP**: Writing – Review & Editing. **MAB**: Methodology, Investigation, Writing – Review & Editing. **MRS**: Conceptualization, Methodology, Writing – Review & Editing, Funding Acquisition, Supervision. **CEM**: Methodology, Writing – Review & Editing, Resources. **RB**: Writing – Review & Editing, Formal Analysis, Validation, Data Curation. **TMB**: Conceptualization, Methodology, Writing – Review & Editing, Funding Acquisition, Supervision, Project Administration. **TMB** is the guarantor of this work and, as such, has full access to all of the data in the study and takes responsibility for the integrity of the data and the accuracy of the data analysis. All authors contributed to the article and approved the submitted version.

## References

[1] Rodriguez-Calvo T, Johnson JD, Overbergh L, Dunne JL (2021) Neoepitopes in Type 1 Diabetes: Etiological Insights, Biomarkers and Therapeutic Targets. Front Immunol 12: 667989. 10.3389/fimmu.2021.667989

[2] Atkinson MA, Eisenbarth GS, Michels AW (2014) Type 1 diabetes. Lancet 383(9911): 69–82. 10.1016/S0140-6736(13)60591-7

[3] Herold KC, Bundy BN, Long SA, et al. (2019) An Anti-CD3 Antibody, Teplizumab, in Relatives at Risk for Type 1 Diabetes. N Engl J Med 381(7): 603–613. 10.1056/NEJMoa1902226

[4] Herold KC, Gitelman SE, Ehlers MR, et al. (2013) Teplizumab (anti-CD3 mAb) treatment preserves C-peptide responses in patients with new-onset type 1 diabetes in a randomized controlled trial: metabolic and immunologic features at baseline identify a subgroup of responders. Diabetes 62(11): 3766–3774. 10.2337/db13-0345

[5] Loftus EV, Panés J, Lacerda AP, et al. (2023) Upadacitinib Induction and Maintenance Therapy for Crohn’s Disease. N Engl J Med 388(21): 1966–1980. 10.1056/NEJMoa2212728

[6] Korhonen R, Moilanen E (2010) Anti-CD20 antibody rituximab in the treatment of rheumatoid arthritis. Basic Clin Pharmacol Toxicol 106(1): 13–21. 10.1111/j.1742-7843.2009.00452.x

[7] Ayano M, Tsukamoto H, Kohno K, et al. (2015) Increased CD226 Expression on CD8+ T Cells Is Associated with Upregulated Cytokine Production and Endothelial Cell Injury in Patients with Systemic Sclerosis. J Immunol 195(3): 892–900. 10.4049/jimmunol.1403046

[8] Fuhrman CA, Yeh WI, Seay HR, et al. (2015) Divergent Phenotypes of Human Regulatory T Cells Expressing the Receptors TIGIT and CD226. J Immunol 195(1): 145–155. 10.4049/jimmunol.1402381

[9] Hou S, Ge K, Zheng X, Wei H, Sun R, Tian Z (2014) CD226 protein is involved in immune synapse formation and triggers Natural Killer (NK) cell activation via its first extracellular domain. J Biol Chem 289(10): 6969–6977. 10.1074/jbc.M113.498253

[10] Lozano E, Dominguez-Villar M, Kuchroo V, Hafler DA (2012) The TIGIT/CD226 axis regulates human T cell function. J Immunol 188(8): 3869–3875. 10.4049/jimmunol.1103627

[11] Zhang Z, Wu N, Lu Y, Davidson D, Colonna M, Veillette A (2015) DNAM-1 controls NK cell activation via an ITT-like motif. J Exp Med 212(12): 2165–2182. 10.1084/jem.20150792

[12] Lozano E, Joller N, Cao Y, Kuchroo VK, Hafler DA (2013) The CD226/CD155 interaction regulates the proinflammatory (Th1/Th17)/anti-inflammatory (Th2) balance in humans. J Immunol 191(7): 3673–3680. 10.4049/jimmunol.1300945

[13] Liu T, Zhang D, Zhang Y, et al. (2018) Blocking CD226 Promotes Allogeneic Transplant Immune Tolerance and Improves Skin Graft Survival by Increasing the Frequency of Regulatory T Cells in a Murine Model. Cell Physiol Biochem 45(6): 2338–2350. 10.1159/000488182

[14] Qiu ZX, Zhang K, Qiu XS, Zhou M, Li WM (2013) CD226 Gly307Ser association with multiple autoimmune diseases: a meta-analysis. Hum Immunol 74(2): 249–255. 10.1016/j.humimm.2012.10.009

[15] Shapiro MR, Thirawatananond P, Peters L, et al. (2021) De-coding genetic risk variants in type 1 diabetes. Immunol Cell Biol 99(5): 496–508. 10.1111/imcb.12438

[16] Gaud G, Roncagalli R, Chaoui K, et al. (2018) The costimulatory molecule CD226 signals through VAV1 to amplify TCR signals and promote IL-17 production by CD4. Sci Signal 11(538). 10.1126/scisignal.aar3083

[17] Shibuya K, Shirakawa J, Kameyama T, et al. (2003) CD226 (DNAM-1) is involved in lymphocyte function-associated antigen 1 costimulatory signal for naive T cell differentiation and proliferation. J Exp Med 198(12): 1829–1839. 10.1084/jem.20030958

[18] Shirakawa J, Shibuya K, Shibuya A (2005) Requirement of the serine at residue 329 for lipid raft recruitment of DNAM-1 (CD226). Int Immunol 17(3): 217–223. 10.1093/intimm/dxh199

[19] Hollis-Moffatt JE, Hook SM, Merriman TR (2005) Colocalization of mouse autoimmune diabetes loci Idd21.1 and Idd21.2 with IDDM6 (human) and Iddm3 (rat). Diabetes 54(9): 2820–2825. 10.2337/diabetes.54.9.2820

[20] Shapiro MR, Yeh WI, Longfield JR, et al. (2020) CD226 Deletion Reduces Type 1 Diabetes in the NOD Mouse by Impairing Thymocyte Development and Peripheral T Cell Activation. Front Immunol 11: 2180. 10.3389/fimmu.2020.02180

[21] Wang N, Yi H, Fang L, et al. (2020) CD226 Attenuates Treg Proliferation via Akt and Erk Signaling in an EAE Model. Front Immunol 11: 1883. 10.3389/fimmu.2020.01883

[22] Zhang R, Zeng H, Zhang Y, et al. (2016) CD226 ligation protects against EAE by promoting IL-10 expression via regulation of CD4+ T cell differentiation. Oncotarget 7(15): 19251–19264. 10.18632/oncotarget.7834

[23] Thirawatananond P, Brown ME, Sachs LK, et al. (2023) Treg-Specific CD226 Deletion Reduces Diabetes Incidence in NOD Mice by Improving Regulatory T-Cell Stability. Diabetes 72(11): 1629–1640. 10.2337/db23-0307

[24] Zhou X, Jeker LT, Fife BT, Zhu S, Anderson MS, McManus MT, Bluestone JA (2008) Selective miRNA disruption in T reg cells leads to uncontrolled autoimmunity. J Exp Med 205(9): 1983–1991. 10.1084/jem.20080707

[25] Whitener RL, Gallo Knight L, Li J, et al. (2017) The Type 1 Diabetes–Resistance Locus Idd22 Controls Trafficking of Autoreactive CTLs into the Pancreatic Islets of NOD Mice. The Journal of Immunology 199(12). 10.4049/jimmunol.1602037

[26] National Research Council. Committee for the Update of the Guide for the C, Use of Laboratory A, Institute for Laboratory Animal R, National Academies P (2011) Guide for the care and use of laboratory animals. 8th edn. National Academies Press, Washington, D.C.

[27] Xue S, Posgai A, Wasserfall C, et al. (2015) Combination Therapy Reverses Hyperglycemia in NOD Mice With Established Type 1 Diabetes. Diabetes 64(11): 3873–3884. 10.2337/db15-0164

28. Antov A, Yang L, Vig M, Baltimore D, Van Parijs L (2003) Essential role for STAT5 signaling in CD25+CD4+ regulatory T cell homeostasis and the maintenance of self-tolerance. J Immunol 171(7): 3435–3441. 10.4049/jimmunol.171.7.3435

[29] You S, Chen C, Lee WH, Brusko T, Atkinson M, Liu CP (2004) Presence of diabetes-inhibiting, glutamic acid decarboxylase-specific, IL-10-dependent, regulatory T cells in naive nonobese diabetic mice. J Immunol 173(11): 6777-6785. 10.4049/jimmunol.173.11.6777

[30] Lissina A, Ladell K, Skowera A, et al. (2009) Protein kinase inhibitors substantially improve the physical detection of T-cells with peptide-MHC tetramers. J Immunol Methods 340(1): 11–24. 10.1016/j.jim.2008.09.014

[31] Chen J, Grieshaber S, Mathews CE (2011) Methods to assess beta cell death mediated by cytotoxic T lymphocytes. J Vis Exp(52). 10.3791/2724

[32] Lamont D, Mukherjee G, Kumar PR, et al. (2014) Compensatory mechanisms allow undersized anchor-deficient class I MHC ligands to mediate pathogenic autoreactive T cell responses. J Immunol 193(5): 2135–2146. 10.4049/jimmunol.1400997

[33] Hamaguchi K, Gaskins HR, Leiter EH (1991) NIT-1, a pancreatic beta-cell line established from a transgenic NOD/Lt mouse. Diabetes 40(7): 842–849. 10.2337/diab.40.7.842

[34] Dobin A, Davis CA, Schlesinger F, et al. (2013) STAR: ultrafast universal RNA-seq aligner. Bioinformatics 29(1): 15–21. 10.1093/bioinformatics/bts635

[35] Choudhary S, Satija R (2022) Comparison and evaluation of statistical error models for scRNA-seq. Genome Biol 23(1): 27. 10.1186/s13059-021-02584-9

[36] Ahlmann-Eltze C, Huber W (2021) glmGamPoi: fitting Gamma-Poisson generalized linear models on single cell count data. Bioinformatics 36(24): 5701–5702. 10.1093/bioinformatics/btaa1009

[37] Pijuan-Sala B, Griffiths JA, Guibentif C, et al. (2019) A single-cell molecular map of mouse gastrulation and early organogenesis. Nature 566(7745): 490–495. 10.1038/s41586-019-0933-9

[38] Lun ATL, Richard AC, Marioni JC (2017) Testing for differential abundance in mass cytometry data. Nat Methods 14(7): 707–709. 10.1038/nmeth.4295

[39] Kasmani MY, Ciecko AE, Brown AK, Petrova G, Gorski J, Chen YG, Cui W (2022) Autoreactive CD8 T cells in NOD mice exhibit phenotypic heterogeneity but restricted TCR gene usage. Life Sci Alliance 5(10). 10.26508/lsa.202201503

[40] Hao Y, Stuart T, Kowalski MH, et al. (2023) Dictionary learning for integrative, multimodal and scalable single-cell analysis. Nat Biotechnol. 10.1038/s41587-023-01767-y

[41] Mulè MP, Martins AJ, Tsang JS (2022) Normalizing and denoising protein expression data from droplet-based single cell profiling. Nat Commun 13(1): 2099. 10.1038/s41467-022-29356-8

[42] McGinnis CS, Murrow LM, Gartner ZJ (2019) DoubletFinder: Doublet Detection in Single-Cell RNA Sequencing Data Using Artificial Nearest Neighbors. Cell Syst 8(4): 329–337.e324. 10.1016/j.cels.2019.03.003

43. Korsunsky I, Nathan A, Millard N, Raychaudhuri S (2019) Presto scales Wilcoxon and auROC analyses to millions of observations. In, BioRxiv

[44] Chen Y, Lun AT, Smyth GK (2016) From reads to genes to pathways: differential expression analysis of RNA-Seq experiments using Rsubread and the edgeR quasi-likelihood pipeline. F1000Res 5: 1438. 10.12688/f1000research.8987.2

[45] Amezquita RA, Lun ATL, Becht E, et al. (2020) Orchestrating single-cell analysis with Bioconductor. Nat Methods 17(2): 137–145. 10.1038/s41592-019-0654-x

[46] Borcherding N, Bormann NL, Kraus G (2020) scRepertoire: An R-based toolkit for single-cell immune receptor analysis. F1000Res 9: 47. 10.12688/f1000research.22139.2

[47] Pagès H, Aboyoun P, Gentleman R, DebRoy S (2024) Biostrings: Efficient manipulation of biological strings. In. R package version 2.70.2

[48] Bunis DG, Andrews J, Fragiadakis GK, Burt TD, Sirota M (2021) dittoSeq: universal user-friendly single-cell and bulk RNA sequencing visualization toolkit. Bioinformatics 36(22-23): 5535–5536. 10.1093/bioinformatics/btaa1011

[49] Blighe K, Rana S, Lewis M (2023) EnhancedVolcano: Publication-ready volcano plots with enhanced colouring and labeling. In. R package version 1.20.0

[50] Akimova T, Levine MH, Beier UH, Hancock WW (2016) Standardization, Evaluation, and Area-Under-Curve Analysis of Human and Murine Treg Suppressive Function. Methods Mol Biol 1371: 43–78. 10.1007/978-1-4939-3139-2_4

[51] Brown ME, Peters LD, Hanbali SR, et al. (2022) Human CD4+ CD25+ CD226-Tregs Demonstrate Increased Purity, Lineage Stability, and Suppressive Capacity Versus CD4+ CD25+ CD127lo/- Tregs for Adoptive Cell Therapy. Front Immunol 13: 873560. 10.3389/fimmu.2022.873560

[52] Permanyer M, Bošnjak B, Glage S, et al. (2021) Efficient IL-2R signaling differentially affects the stability, function, and composition of the regulatory T-cell pool. Cell Mol Immunol 18(2): 398–414. 10.1038/s41423-020-00599-z

[53] Brusko TM, Russ HA, Stabler CL (2021) Strategies for durable β cell replacement in type 1 diabetes. Science 373(6554): 516–522. 10.1126/science.abh1657

[54] Tahara-Hanaoka S, Shibuya K, Onoda Y, et al. (2004) Functional characterization of DNAM-1 (CD226) interaction with its ligands PVR (CD155) and nectin-2 (PRR-2/CD112). Int Immunol 16(4): 533–538. 10.1093/intimm/dxh059

[55] Banta KL, Xu X, Chitre AS, et al. (2022) Mechanistic convergence of the TIGIT and PD-1 inhibitory pathways necessitates co-blockade to optimize anti-tumor CD8. Immunity 55(3): 512–526.e519. 10.1016/j.immuni.2022.02.005

[56] Anthony DA, Andrews DM, Watt SV, Trapani JA, Smyth MJ (2010) Functional dissection of the granzyme family: cell death and inflammation. Immunol Rev 235(1): 73–92. 10.1111/j.0105-2896.2010.00907.x

[57] Tamura R, Yoshihara K, Nakaoka H, et al. (2020) XCL1 expression correlates with CD8-positive T cells infiltration and PD-L1 expression in squamous cell carcinoma arising from mature cystic teratoma of the ovary. Oncogene 39(17): 3541–3554. 10.1038/s41388-020-1237-0

[58] Seay HR, Yusko E, Rothweiler SJ, et al. (2016) Tissue distribution and clonal diversity of the T and B cell repertoire in type 1 diabetes. JCI Insight 1(20): e88242. 10.1172/jci.insight.88242

[59] Chiou J, Geusz RJ, Okino ML, et al. (2021) Interpreting type 1 diabetes risk with genetics and single-cell epigenomics. Nature 594(7863): 398–402. 10.1038/s41586-021-03552-w

[60] Spanier JA, Sahli NL, Wilson JC, et al. (2017) Increased Effector Memory Insulin-Specific CD4+ T Cells Correlate with Insulin Autoantibodies in Patients With Recent-Onset Type 1 Diabetes. Diabetes 66(12): 3051–3060. 10.2337/db17-0666

[61] Knörck A, Schäfer G, Alansary D, Richter J, Thurner L, Hoth M, Schwarz EC (2022) Cytotoxic Efficiency of Human CD8+ T Cell Memory Subtypes. Front Immunol 13: 838484. 10.3389/fimmu.2022.838484

[62] Gilfillan S, Chan CJ, Cella M, et al. (2008) DNAM-1 promotes activation of cytotoxic lymphocytes by nonprofessional antigen-presenting cells and tumors. J Exp Med 205(13): 2965–2973. 10.1084/jem.20081752

[63] Magnuson AM, Thurber GM, Kohler RH, Weissleder R, Mathis D, Benoist C (2015) Population dynamics of islet-infiltrating cells in autoimmune diabetes. Proc Natl Acad Sci U S A 112(5): 1511–1516. 10.1073/pnas.1423769112

[64] Basu R, Whitlock BM, Husson J, et al. (2016) Cytotoxic T Cells Use Mechanical Force to Potentiate Target Cell Killing. Cell 165(1): 100–110. 10.1016/j.cell.2016.01.021

[65] Kunimura K, Uruno T, Fukui Y (2020) DOCK family proteins: key players in immune surveillance mechanisms. Int Immunol 32(1): 5–15. 10.1093/intimm/dxz067

[66] Kuzumi A, Yoshizaki A, Matsuda KM, et al. (2021) Interleukin-31 promotes fibrosis and T helper 2 polarization in systemic sclerosis. Nat Commun 12(1): 5947. 10.1038/s41467-021-26099-w

[67] Aboumrad E, Madec AM, Thivolet C (2007) The CXCR4/CXCL12 (SDF-1) signalling pathway protects non-obese diabetic mouse from autoimmune diabetes. Clin Exp Immunol 148(3): 432–439. 10.1111/j.1365-2249.2007.03370.x

68. Walsh DA, Borges da Silva H, Beura LK, Peng C, Hamilton SE, Masopust D, Jameson SC (2019) The Functional Requirement for CD69 in Establishment of Resident Memory CD8. J Immunol 203(4): 946–955. 10.4049/jimmunol.1900052

[69] Kojima H, Kanada H, Shimizu S, et al. (2003) CD226 mediates platelet and megakaryocytic cell adhesion to vascular endothelial cells. J Biol Chem 278(38): 36748–36753. 10.1074/jbc.M300702200

[70] Chen L, Xie X, Zhang X, Jia W, Jian J, Song C, Jin B (2003) The expression, regulation and adhesion function of a novel CD molecule, CD226, on human endothelial cells. Life Sci 73(18): 2373–2382. 10.1016/s0024-3205(03)00606-4

[71] Johnston RJ, Comps-Agrar L, Hackney J, et al. (2014) The immunoreceptor TIGIT regulates antitumor and antiviral CD8(+) T cell effector function. Cancer Cell 26(6): 923–937. 10.1016/j.ccell.2014.10.018

[72] Bancerek J, Poss ZC, Steinparzer I, et al. (2013) CDK8 kinase phosphorylates transcription factor STAT1 to selectively regulate the interferon response. Immunity 38(2): 250–262. 10.1016/j.immuni.2012.10.017

[73] Fuchs YF, Eugster A, Dietz S, et al. (2017) CD8+ T cells specific for the islet autoantigen IGRP are restricted in their T cell receptor chain usage. Sci Rep 7: 44661. 10.1038/srep44661

[74] Unger WW, Pearson T, Abreu JR, et al. (2012) Islet-specific CTL cloned from a type 1 diabetes patient cause beta-cell destruction after engraftment into HLA-A2 transgenic NOD/scid/IL2RG null mice. PLoS One 7(11): e49213. 10.1371/journal.pone.0049213

[75] Ke Q, Kroger CJ, Clark M, Tisch RM (2020) Evolving Antibody Therapies for the Treatment of Type 1 Diabetes. Front Immunol 11: 624568. 10.3389/fimmu.2020.624568

[76] Garo LP, Ajay AK, Fujiwara M, et al. (2019) Smad7 Controls Immunoregulatory PDL2/1-PD1 Signaling in Intestinal Inflammation and Autoimmunity. Cell Rep 28(13): 3353–3366.e3355. 10.1016/j.celrep.2019.07.065

[77] Yan X, Liu Z, Chen Y (2009) Regulation of TGF-beta signaling by Smad7. Acta Biochim Biophys Sin (Shanghai) 41(4): 263–272. 10.1093/abbs/gmp018

[78] Zhou G, Sun X, Qin Q, et al. (2018) Loss of Smad7 Promotes Inflammation in Rheumatoid Arthritis. Front Immunol 9: 2537. 10.3389/fimmu.2018.02537

[79] Pang N, Zhang F, Ma X, et al. (2014) TGF-β/Smad signaling pathway regulates Th17/Treg balance during Echinococcus multilocularis infection. Int Immunopharmacol 20(1): 248–257. 10.1016/j.intimp.2014.02.038

80. Du X, de Almeida P, Manieri N, et al. (2018) CD226 regulates natural killer cell antitumor responses via phosphorylation-mediated inactivation of transcription factor FOXO1. Proc Natl Acad Sci U S A 115(50): E11731–E11740. 10.1073/pnas.1814052115

[81] Ouyang W, Beckett O, Ma Q, Paik JH, DePinho RA, Li MO (2010) Foxo proteins cooperatively control the differentiation of Foxp3+ regulatory T cells. Nat Immunol 11(7): 618–627. 10.1038/ni.1884

[82] Toomer KH, Lui JB, Altman NH, Ban Y, Chen X, Malek TR (2019) Essential and non-overlapping IL-2Rα-dependent processes for thymic development and peripheral homeostasis of regulatory T cells. Nat Commun 10(1): 1037. 10.1038/s41467-019-08960-1

[83] Passerini L, Allan SE, Battaglia M, et al. (2008) STAT5-signaling cytokines regulate the expression of FOXP3 in CD4+CD25+ regulatory T cells and CD4+CD25-effector T cells. Int Immunol 20(3): 421–431. 10.1093/intimm/dxn002

[84] Hull CM, Peakman M, Tree TIM (2017) Regulatory T cell dysfunction in type 1 diabetes: what’s broken and how can we fix it? Diabetologia 60(10): 1839–1850. 10.1007/s00125-017-4377-1

[85] Buckner JH (2010) Mechanisms of impaired regulation by CD4(+)CD25(+)FOXP3(+) regulatory T cells in human autoimmune diseases. Nat Rev Immunol 10(12): 849–859. 10.1038/nri2889

[86] Hafeez U, Gan HK, Scott AM (2018) Monoclonal antibodies as immunomodulatory therapy against cancer and autoimmune diseases. Curr Opin Pharmacol 41: 114–121. 10.1016/j.coph.2018.05.010

[87] Mohty M (2007) Mechanisms of action of antithymocyte globulin: T-cell depletion and beyond. Leukemia 21(7): 1387–1394. 10.1038/sj.leu.2404683

[88] Haller MJ, Schatz DA, Skyler JS, et al. (2018) Low-Dose Anti-Thymocyte Globulin (ATG) Preserves β-Cell Function and Improves HBA1c, and Increases Regulatory to Conventional T-Cell Ratios in New-Onset Type 1 Diabetes: Two-Year Clinical Trial Data. Diabetes Care 41(9): 1917–1925. 10.2337/dc18-0494

[89] Jacobsen LM, Diggins K, Blanchfield L, et al. (2023) Responders to low-dose ATG induce CD4+ T cell exhaustion in type 1 diabetes. JCI Insight 8(16). 10.1172/jci.insight.161812

[90] MacIsaac J, Siddiqui R, Jamula E, et al. (2018) Systematic review of rituximab for autoimmune diseases: a potential alternative to intravenous immune globulin. Transfusion 58(11): 2729–2735. 10.1111/trf.14841

[91] Long SA, Thorpe J, DeBerg HA, et al. (2016) Partial exhaustion of CD8 T cells and clinical response to teplizumab in new-onset type 1 diabetes. Sci Immunol 1(5). 10.1126/sciimmunol.aai7793

[92] Long SA, Thorpe J, Herold KC, et al. (2017) Remodeling T cell compartments during anti-CD3 immunotherapy of type 1 diabetes. Cell Immunol 319: 3–9. 10.1016/j.cellimm.2017.07.007

[93] Burns GF, Triglia T, Werkmeister JA, Begley CG, Boyd AW (1985) TLiSA1, a human T lineage-specific activation antigen involved in the differentiation of cytotoxic T lymphocytes and anomalous killer cells from their precursors. J Exp Med 161(5): 1063–1078. 10.1084/jem.161.5.1063

[94] Scott JL, Dunn SM, Jin B, et al. (1989) Characterization of a novel membrane glycoprotein involved in platelet activation. J Biol Chem 264(23): 13475–13482

[95] Reymond N, Imbert AM, Devilard E, et al. (2004) DNAM-1 and PVR regulate monocyte migration through endothelial junctions. J Exp Med 199(10): 1331–1341. 10.1084/jem.20032206

[96] Ma D, Sun Y, Lin D, et al. (2005) CD226 is expressed on the megakaryocytic lineage from hematopoietic stem cells/progenitor cells and involved in its polyploidization. Eur J Haematol 74(3): 228–240. 10.1111/j.1600-0609.2004.00345.x

[97] Aldrich VR, Hernandez-Rovira BB, Chandwani A, Abdulreda MH (2020) NOD Mice-Good Model for T1D but Not Without Limitations. Cell Transplant 29: 963689720939127. 10.1177/0963689720939127

