## Supplemental Materials for "Inhibition of CD226 Co-Stimulation Suppresses Diabetes Development in the NOD Mouse by Augmenting Tregs and Diminishing Effector T Cell Function"

**Supplemental Tables:**

**Supplemental Table 1. Antibodies used for murine flow cytometry.**

| **Target** | **Clone** | **Host** | **Fluorochrome** | **Concentration** | **Vendor** | **RRID** |
| --- | --- | --- | --- | --- | --- | --- |
| CD4 | GK1.5 | Rat | PerCP-Cy5.5 | 0.2 µg/mL | BioLegend | AB_893324 |
| CD4 | RM4-5 | Rat | PE-Cy7 | 0.2 µg/mL | BioLegend | AB_312729 |
| CD8α | 53-6.7 | Rat | BV711 | 0.06 µg/mL | BioLegend | AB_2562100 |
| CD16/CD32 | 2.4G2 | Rat | N/A | 2 µg/mL | BD Pharmingen | AB_394656 |
| CD25 | PC61 | Rat | APC | 0.4 µg/mL | BioLegend | AB_493709 |
| CD44 | IM7 | Rat | PE | 0.4 µg/mL | BioLegend | AB_312959 |
| CD62L | MEL-14 | Rat | APC | 0.4 µg/mL | BioLegend | AB_313099 |
| CD226 | TX42.1 | Rat | BV650 | 0.4 µg/mL | BioLegend | AB_2716083 |
| Foxp3 | FJK-16s | Rat | AF488 | 0.8 µg/mL | eBioScience | AB_763537 |
| Helios | 22F6 | Armenian Hamster | Pacific Blue | 0.8 µg/mL | BioLegend | AB_10690535 |
| Ki-67 | 16A8 | Rat | PE-Cy7 | 0.8 µg/mL | BioLegend | AB_2632694 |
| NKp46 | 29A1.4 | Rat | AF647 | 0.4 µg/mL | BioLegend | AB_2566360 |
| pSTAT5 | C71E5 | Rabbit | AF647 | 0.4 µg/mL | Cell Signaling | AB_2302702 |
| Rat IgG2a | MRG2a-83 | Mouse | FITC | 0.4 µg/mL | BioLegend | AB_492924 |

**Supplemental Table 2. Antibodies used for CITE-seq Staining.**

| **Target** | **Clone** | **Host** | **Barcode** | **Concentration** | **Vendor** | **RRID** |
| --- | --- | --- | --- | --- | --- | --- |
| CD4 | RM4-5 | Rat | AACAAGACCCTTGAG | 10 µg/mL | BioLegend | AB_2813913 |
| CD8α | 53-6.7 | Rat | TACCCGTAATAGCGT | 10 µg/mL | BioLegend | AB_2819776 |
| CD16/CD32 | S17011E | Rat | N/A | 12.5 µg/mL | BioLegend | AB_2783138 |
| CD44 | IM7 | Rat | TGGCTTCAGGTCCTA | 10 µg/mL | BioLegend | AB_2813930 |
| CD62L | MEL-14 | Rat | TGGGCCTAAGTCATC | 10 µg/mL | BioLegend | AB_2819800 |
| TIGIT | 1G9 | Mouse | GAAAGTCGCCAACAG | 10 µg/mL | BioLegend | AB_2819886 |

**Supplemental Figures:**

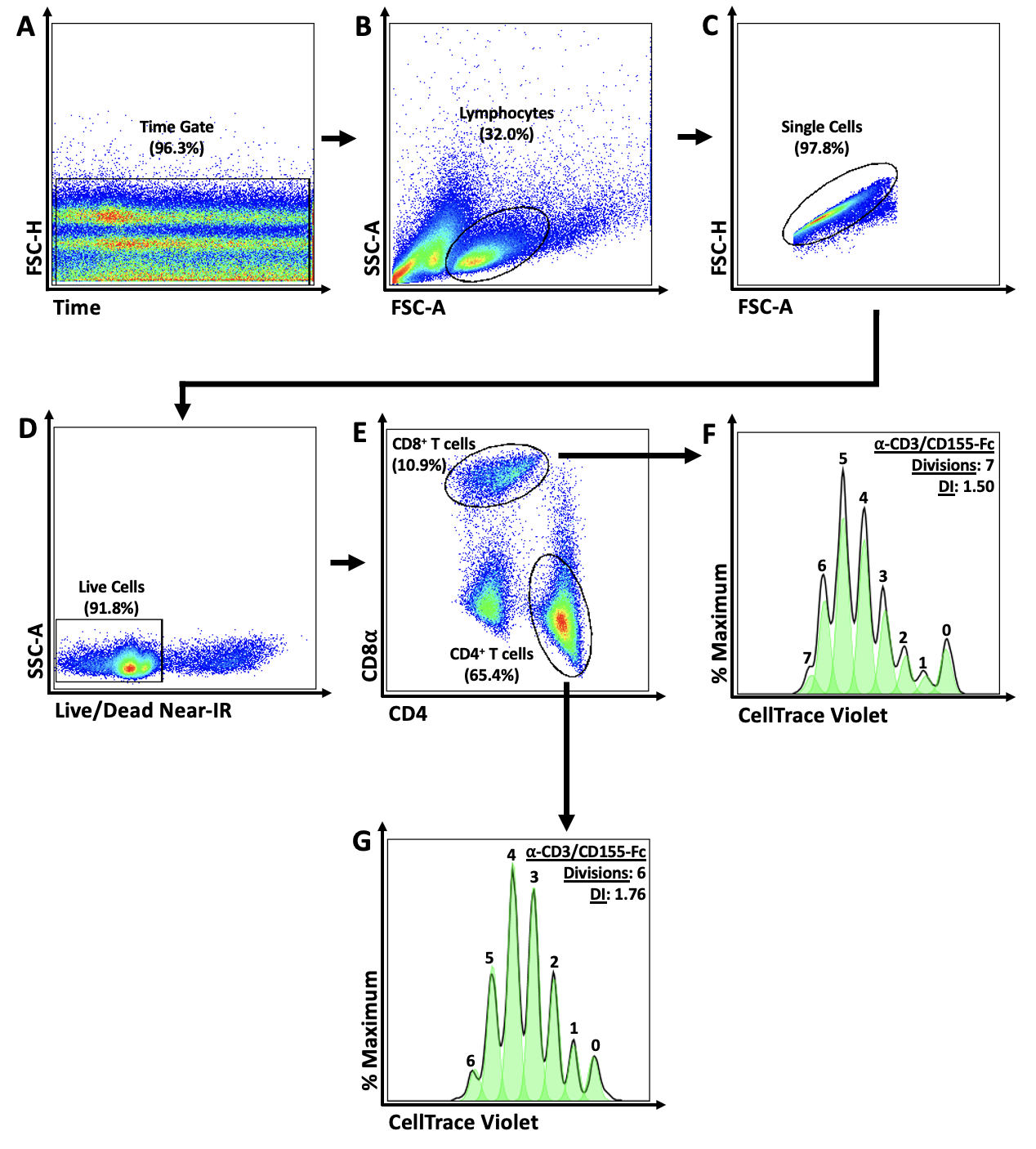

**Supplemental Figure 1.** **Gating Strategy for *In Vitro* Proliferation Assays**. Representative flow plots demonstrate how splenic CD4^+^ T cells and CD8^+^ T cells were identified for evaluating *in vitro* proliferation following treatment with isotype control or anti-CD226 mAb using a Cytek Aurora 5L Spectral Flow Cytometer. (**A**) Cells collected within a stable flow stream were gated using a time by forward scatter height (FSC-H) plot. (**B**) Lymphocyte gating was accomplished using forward scatter area (FSC-A) by side scatter area (SSC-A). (**C**) Single cells were gated upon using FSC-A by FSC-H. (**D**) Live lymphocytes were identified using a Live/Dead Near-IR dye exclusion. (**E**) Single-positive CD4^+^ and CD8^+^ T cell subsets within the live population were gated, before evaluating CellTrace Violet dye dilution was used to assess Tresp proliferation after 4 days of co-culture with Tregs.

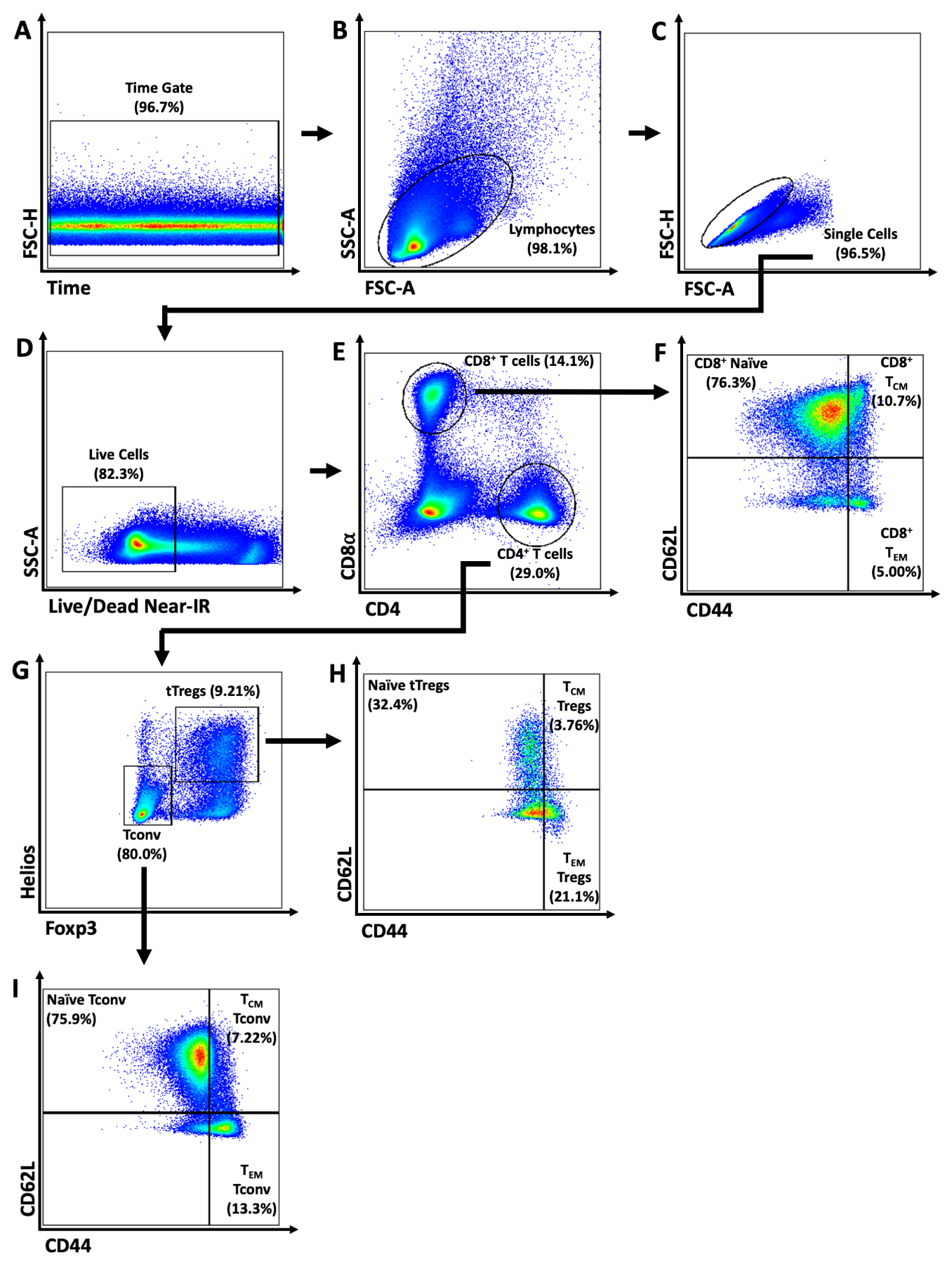
**Supplemental Figure 2. Gating Strategy for tTreg and Memory T cell Subsets**. Representative flow plots demonstrate how thymic Tregs (tTregs) and T cell memory subsets were defined for T cell immunophenotyping following treatment with isotype control or anti-CD226 mAb using a Cytek Aurora 5L Spectral Flow Cytometer. (**A**) Cells collected within a stable flow stream were gated using a time by forward scatter height (FSC-H) plot. (**B**) Lymphocyte gating was accomplished using forward scatter area (FSC-A) by side scatter area (SSC-A). (**C**) Single cells were gated upon using FSC-A by FSC-H. (**D**) Live lymphocytes were identified using a Live/Dead Near-IR dye exclusion to dying cells. (**E**) Single-positive CD4^+^ and CD8^+^ T cell subsets within the live population were gated before defining (**F**) CD8^+^ Naïve (CD44^-^CD62L^+^), T central Memory (T_CM_; CD44^+^CD62L^+^), and T effector Memory (TEM; CD44^+^CD62L^-^) cells using a CD44 by CD62L plot. (**G**) CD4^+^ T cells were further defined as Foxp3^-^Helios^-^ Tconv or Foxp3^+^Helios^+^ tTregs, before identifying (**H**) CD4^+^ tTreg and (**I**) Tconv memory subsets using CD44 by CD62L plots, as previously described.

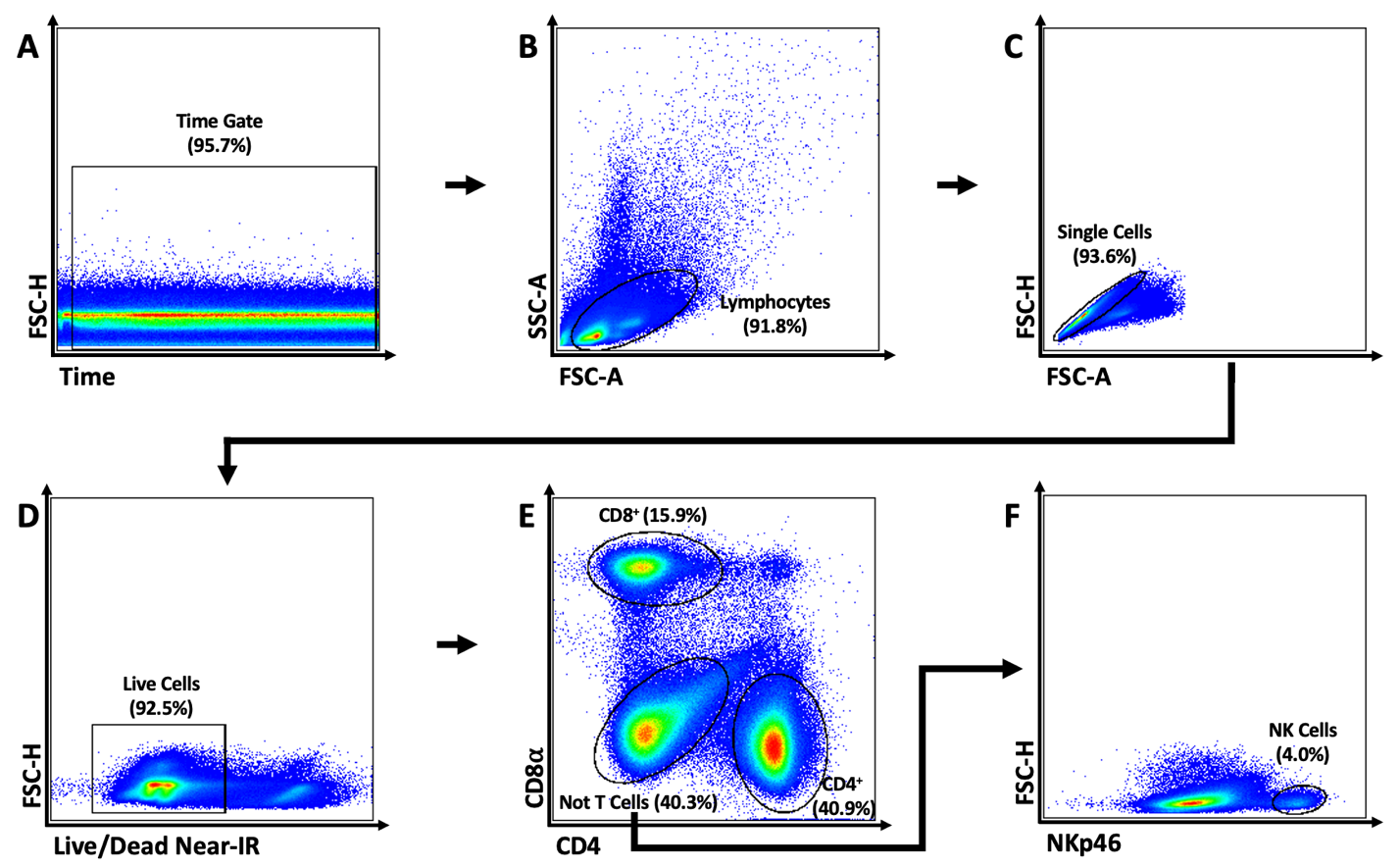
**Supplemental Figure 3.** **Gating Strategy for Detection of anti-CD226 (Clone 480.1) Blocking Antibody**. Representative flow plots demonstrate how splenic CD4^+^ T cells, CD8^+^ T cells, and natural killer (NK) cells were defined for evaluating *in vivo* persistence of anti-CD226 mAb using a Cytek Aurora 5L Spectral Flow Cytometer. (**A**) Cells collected within a stable flow stream were gated using a time by forward scatter height (FSC-H) plot. (**B**) Lymphocyte gating was accomplished using forward scatter area (FSC-A) by side scatter area (SSC-A). (**C**) Single cells were gated upon using FSC-A by FSC-H. (**D**) Live lymphocytes were identified using a Live/Dead Near-IR dye exclusion. (**E**) Single-positive CD4^+^ and CD8^+^ T cell subsets within the live population were gated, whereas (**F**) NK receptor-expressing cells were defined as CD4^-^CD8^-^ lymphocytes expressing NKp46.

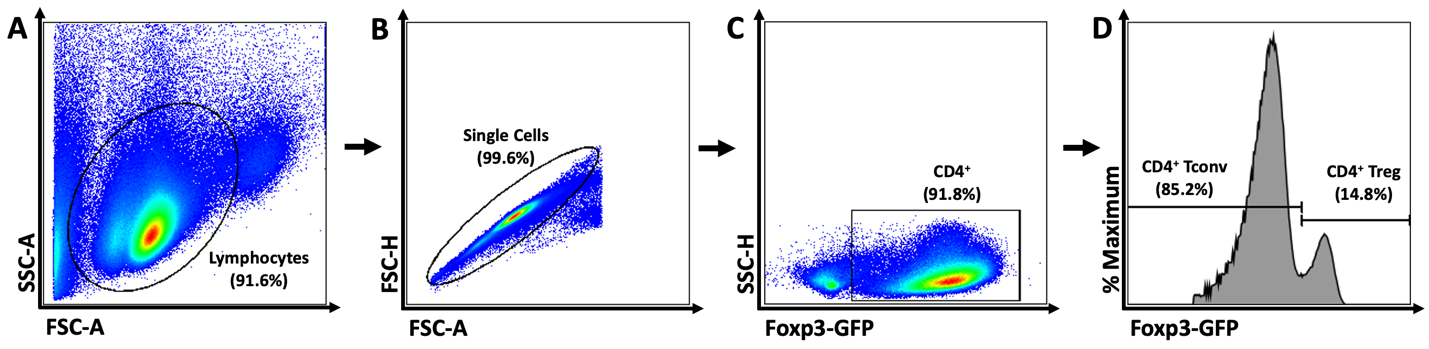
**Supplemental Figure 4.** **Fluorescence Activated Cell Sorting (FACS) Isolation Strategy for CD4^+^ Tconv and Tregs**. Representative flow plots demonstrate how splenic CD4^+^ Tconv and Tregs were isolated for *in vitro* suppression assays using a BD FACSMelody Cell Sorter. (**A**) Lymphocyte gating was accomplished using forward scatter area (FSC-A) by side scatter area (SSC-A). (**B**) Single cells were gated upon using FSC-A by forward scatter height (FSC-H). (**C**) CD4^+^ T cells were gated before defining (**D**) GFP^+^ Tregs and GFP^-^ Tconv using the Foxp3-GFP reporter.

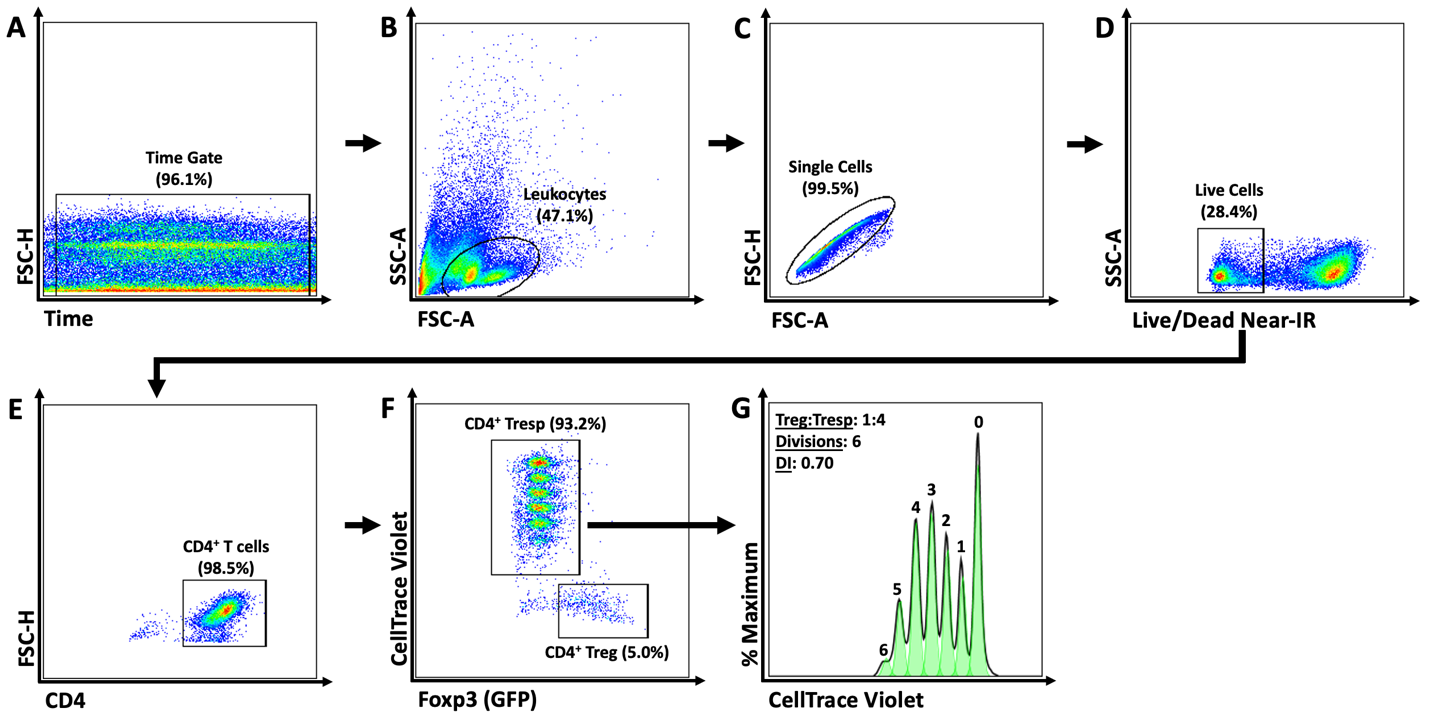
**Supplemental Figure 5.** **Gating Strategy for Assessing *In Vitro* Suppression of CD4^+^ Tresp**. Representative flow plots demonstrate how splenic CD4^+^ T cells, CD8^+^ T cells, and natural killer (NK) cells were defined for evaluating *in vivo* persistence of anti-CD226 mAb using a Cytek Aurora 5L Spectral Flow Cytometer. (**A**) Cells collected within a stable flow stream were gated using a time by forward scatter height (FSC-H) plot. (**B**) Leukocyte gating was accomplished using forward scatter area (FSC-A) by side scatter area (SSC-A). (**C**) Single cells were gated using FSC-A by FSC-H. (**D**) Live leukocytes were identified using a Live/Dead Near-IR dye exclusion. (**E**) CD4^+^ T cells within the live population were gated, before defining (**F**) CellTrace Violet (CTV)^-^GFP^+^ Tregs and CTV^+^GFP^-^ Tresp. (**G**) CTV dye dilution was used to assess Tresp proliferation after 4 days of co-culture with Tregs.

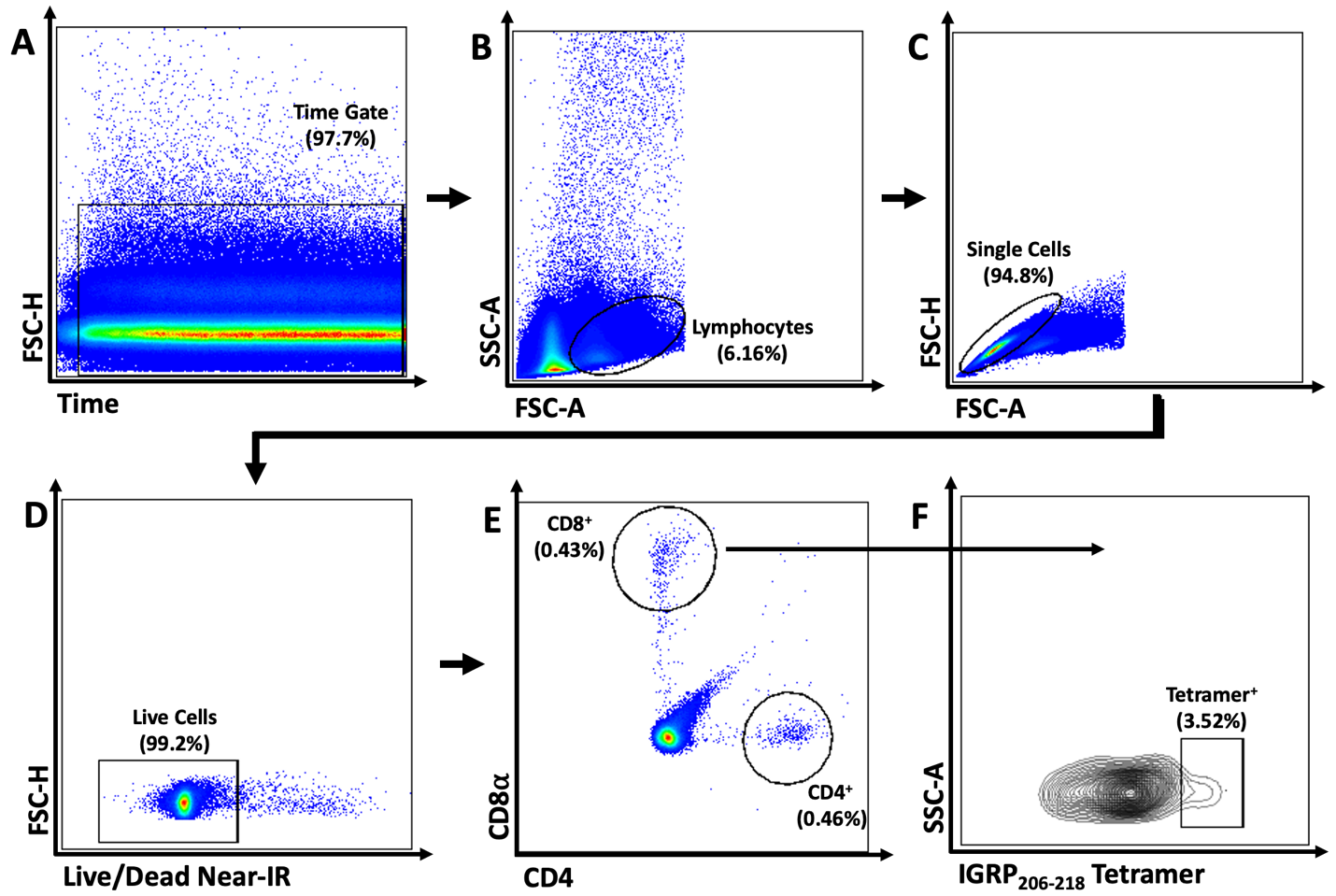
**Supplemental Figure 6.** **Gating Strategy for IGRP_206-218_ Tetramer Staining**. Representative flow plots demonstrate how IGRP-reactive pancreatic CD8^+^ T cells were defined using a Cytek Aurora 5L Spectral Flow Cytometer. (**A**) Cells collected within a stable flow stream were gated using a time by forward scatter height (FSC-H) plot. (**B**) Lymphocyte gating was accomplished using forward scatter area (FSC-A) by side scatter area (SSC-A). (**C**) Single cells were gated upon using FSC-A by FSC-H. (**D**) Live lymphocytes were identified using a Live/Dead Near-IR dye exclusion. (**E**) Single-positive CD4^+^ and CD8^+^ T cell subsets within the live population were gated, before defining (**F**) IGRP_206-218_ tetramer-positive CD8^+^ T cells.

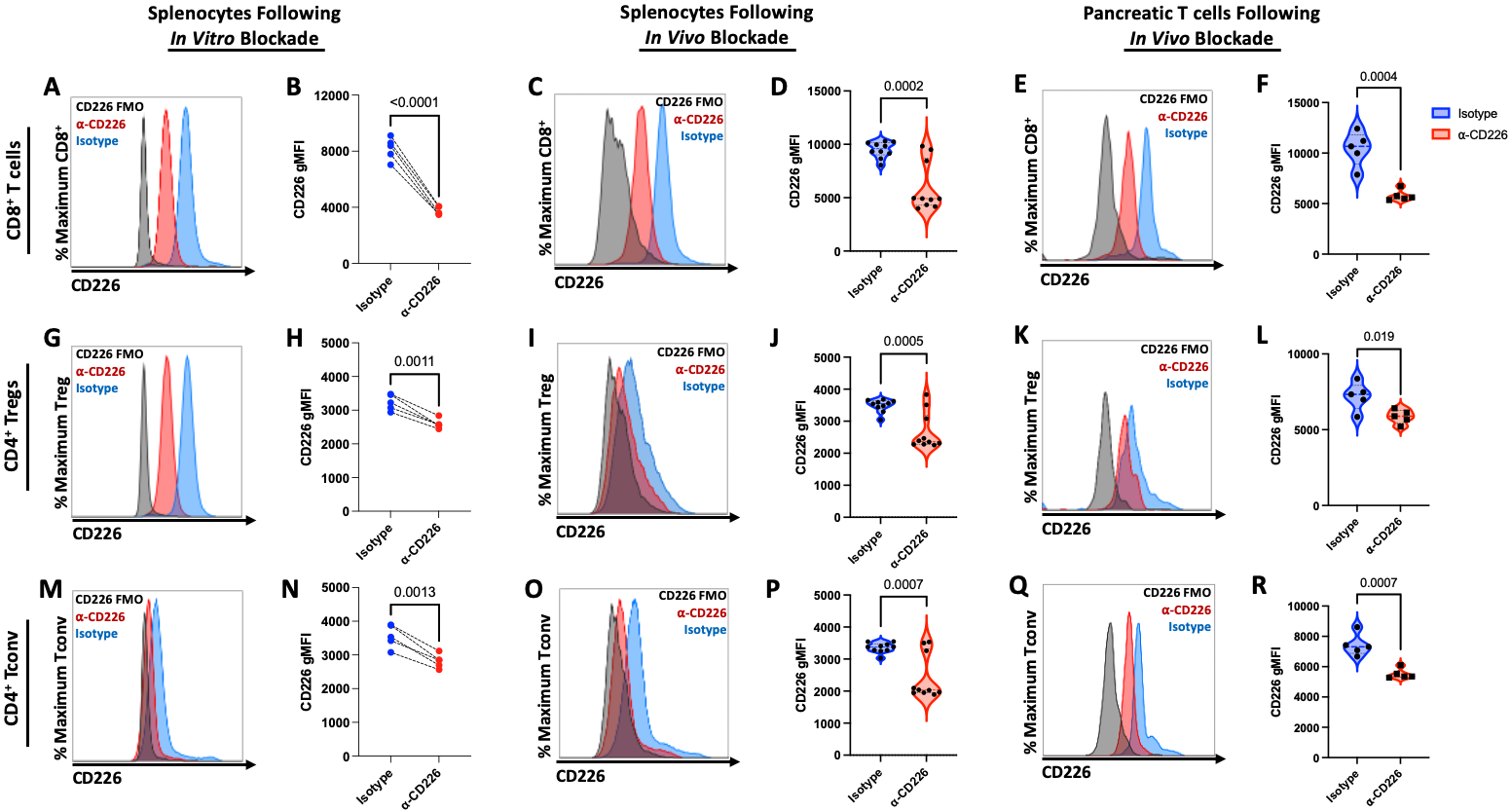
**Supplemental Figure 7. CD226 Accessibility is Reduced Following anti-CD226 mAb Treatment**. The accessibility of CD226 on CD8^+^ T cells (**A-F**), CD4^+^ Tregs (**G-L**), and CD4^+^ Tconv (**M-R**) obtained from NOD spleens or pancreases was assessed using flow cytometry following either *in vitro* (**A-B, G-H, M-N,** biological n=5) or *in vivo* (spleen: **C-D, I-J, O-P,** biological n=10; pancreas: **E-F, K-L, Q-R,** biological n=5) treatment with either IgG2a isotype control (blue) or anti-CD226 clone 480.1 (red). Representative histograms show staining with BV650-conjugated anti-CD226 clone TX42.1 (**A, C, E, G, I, K, M, O, Q**) relative to CD226 fluorescence minus one (FMO) controls (black), with paired dot or violin plots showing the gMFI of CD226 (**B, D, F, H, J, L, N, P, R**). Significant P-values are reported on the figure for t-tests of paired *in vitro* samples and unpaired *ex vivo* samples between treatment conditions. See also Supplemental Figure 2.

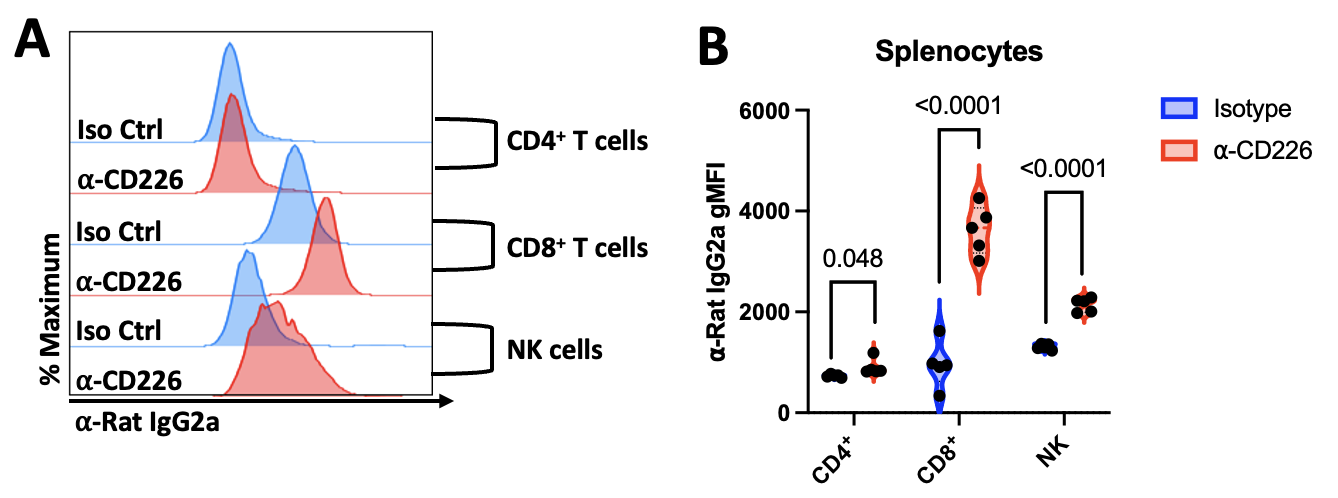
**Supplemental Figure 8.** **Anti-CD226 mAb Persist *In Vivo* for At Least Five Weeks**. Five weeks following *in vivo* treatment with anti-CD226 clone 480.1 mAb (red) or rat IgG2a isotype control (blue), splenocytes were stained with a FITC-labeled ⍺-rat IgG2a mAb to detect cell-bound anti-CD226 or isotype control on three subsets known to express CD226: CD4^+^ T cells, CD8^+^ T cells, and NK cells, as shown in (**A**) Representative histograms and (**B**) violin plots (biological n=5/condition). Significant P-values are reported on the figure for unpaired t-tests for each cell subset between treatment conditions. See also Supplemental Figure 3.

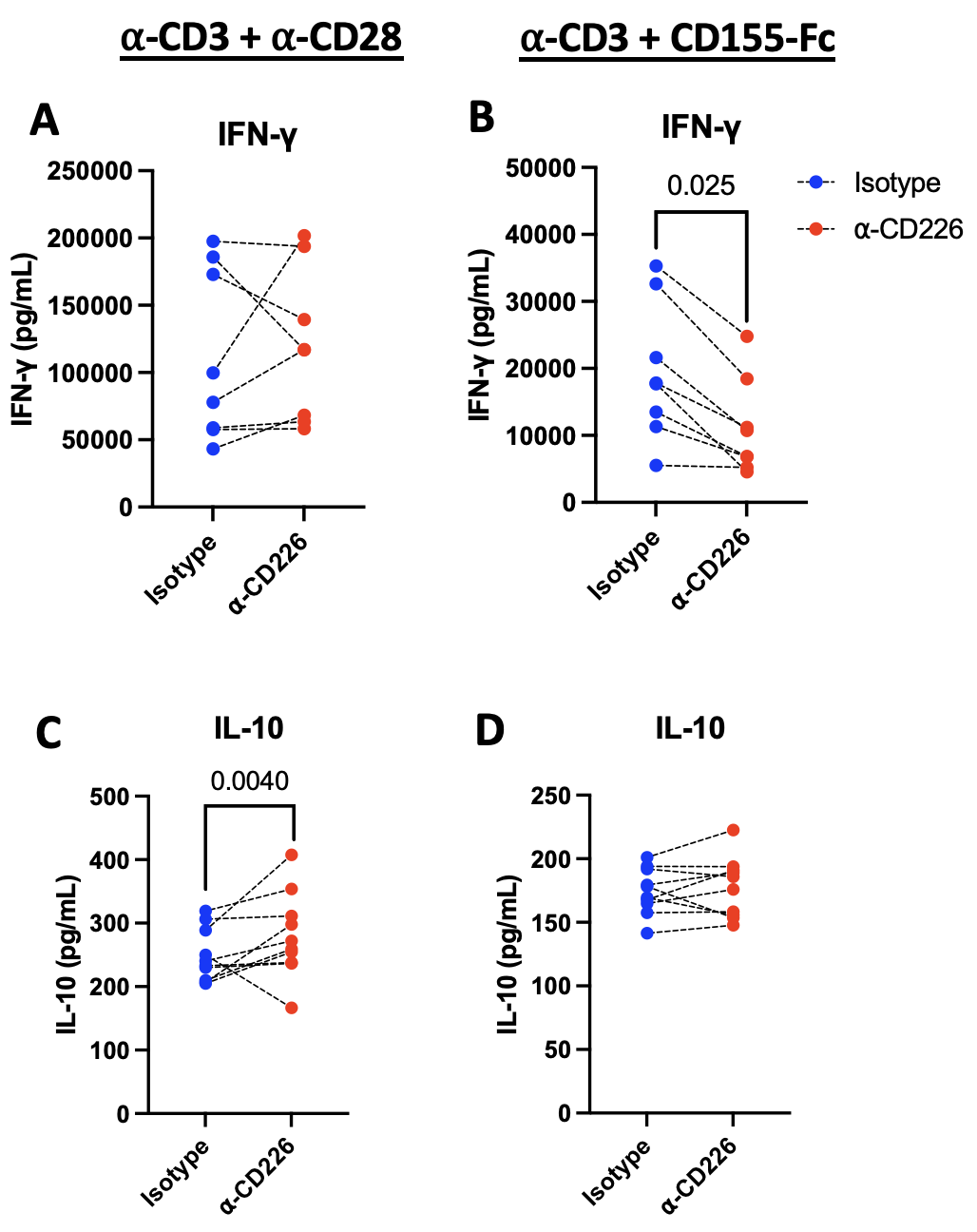

**Supplementary Figure 9. Anti-CD226 mAb Blockade Alters T Cell Cytokine Production**. To assess the impact of anti-CD226 mAb blockade on (**A-B**) IFN-γ and (**C-D**) IL-10 cytokine secretion, cell culture supernatants from splenocytes incubated with IgG2a isotype control (blue) or anti-CD226 mAb (red), then stimulated *in vitro* with either (**A, C**) ⍺-CD3/⍺-CD28 or (**B, D**) ⍺-CD3/CD155-Fc were analyzed by ELISA. Data reflects biological n=8/condition. Paired dot plots show significant P-values reported for two-way ANOVA with Bonferroni correction for multiple comparisons.

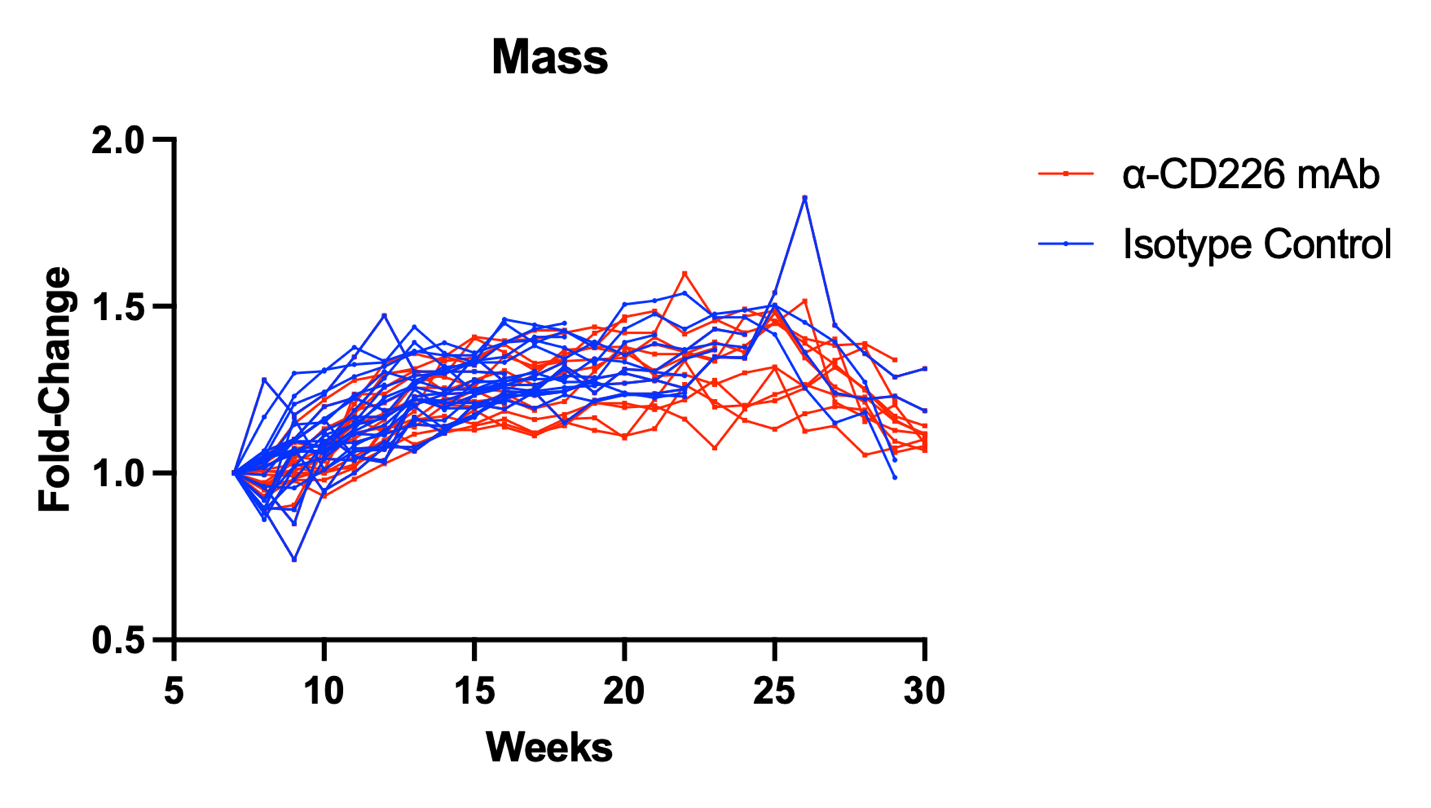
**Supplemental Figure 10. Treatment of NOD mice with anti-CD226 mAb Does Not Impact Body Mass**. Female NOD mice were treated with either IgG2a isotype control (blue) or anti-CD226 mAb (red) and monitored weekly until diabetes onset or 30 weeks of age. Spaghetti plots show fold change in weight from pre-treatment mass at 7 weeks of age through the study endpoint. Data reflects biological n=2-25/treatment group at each time point. No significant P-values were identified for changes in mass using Mixed Effects Analysis with Bonferroni correction for multiple comparisons between treatment conditions.

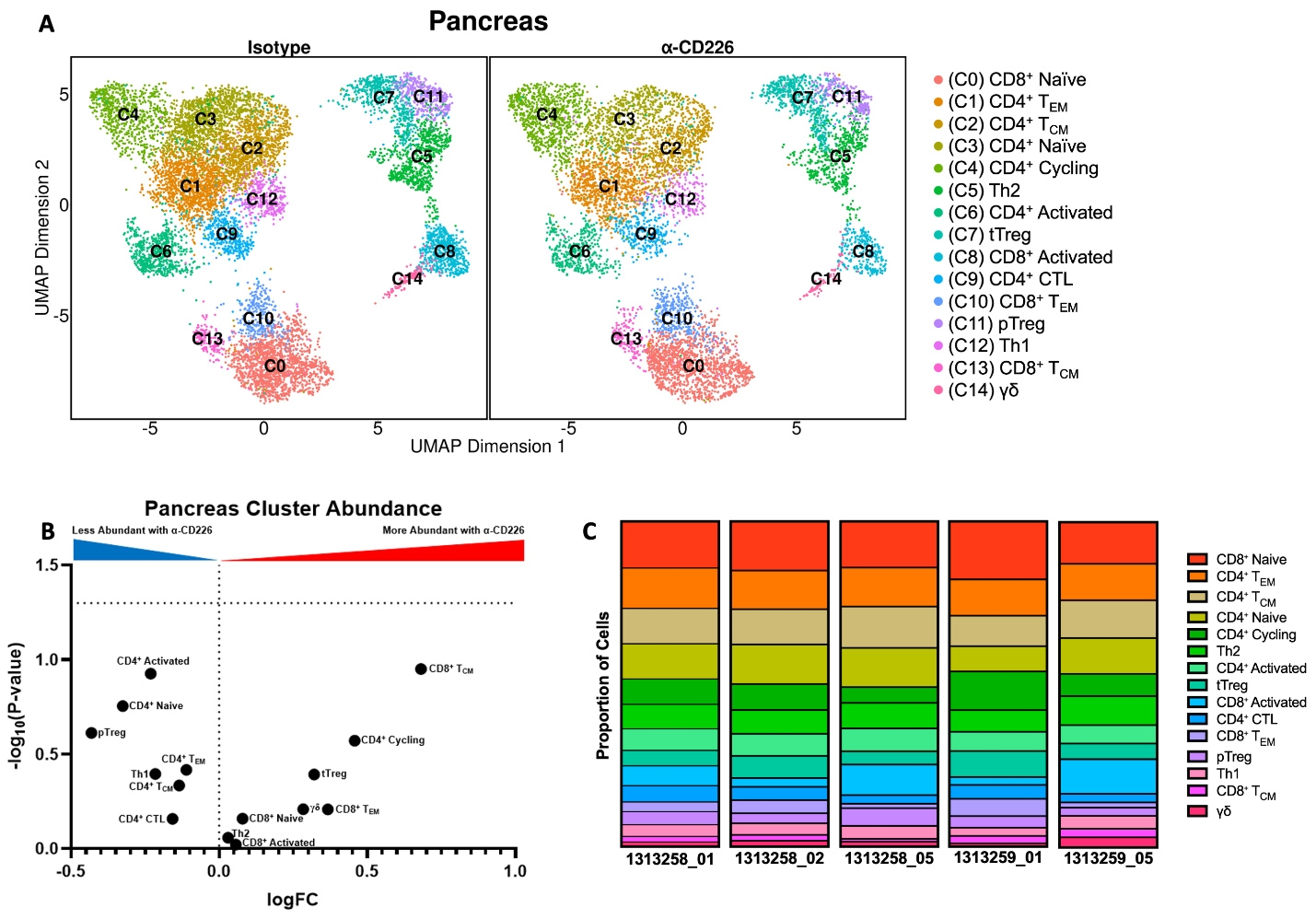
**Supplementary Figure 11. Anti-CD226 mAb Does Not Impact Pancreatic T cell Cluster Abundance**. A differential abundance analysis was performed on T cell clusters identified in the pancreas of 12-week-old female NOD mice 5 weeks after treatment with anti-CD226 mAb or isotype control. (**A**) UMAP projections show the abundance of annotated T cell clusters in the pancreas. (**B**) The volcano plot shows differential abundance values for each T cell cluster with the -log10(P-value) determined by gene-wise negative binomial generalized linear models. (**C**) Stacked bar plots show the proportion of each subset, with each T cell cluster represented similarly between samples.

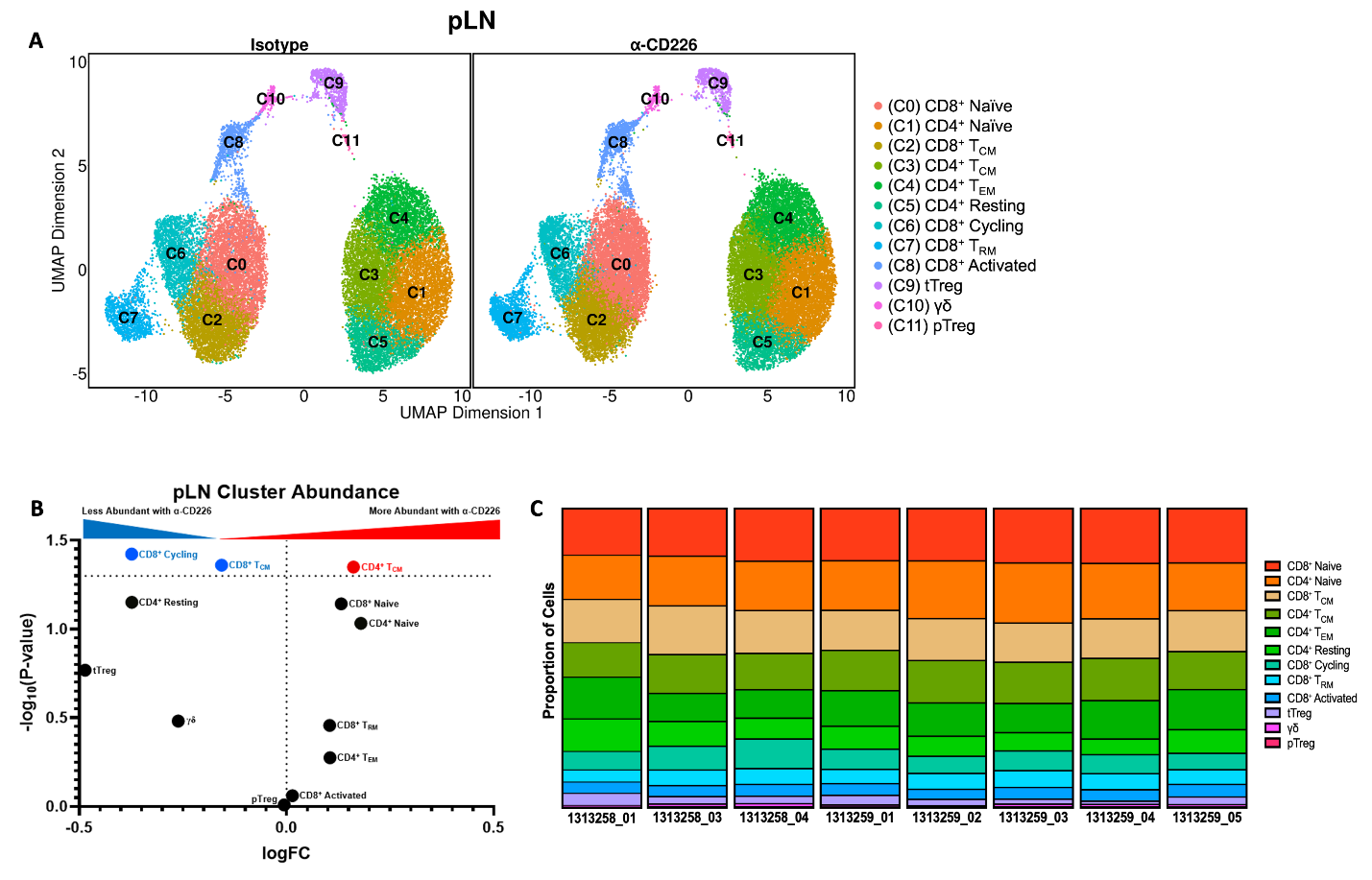
**Supplementary Figure 12. Anti-CD226 mAb Blockade May Alter T Cell Priming and Memory Differentiation in the Pancreatic Lymph Node (pLN)**. A differential abundance analysis was performed on T cell clusters identified in the pLN from 12-week-old female NOD mice 5 weeks after treatment with anti-CD226 mAb or isotype control. (**A**) UMAP projections show the abundance of annotated T cell clusters in pLN. (**B**) The volcano plot shows differential abundance values for each T cell cluster with the -log10(P-value) determined by gene-wise negative binomial generalized linear models. (**C**) Stacked bar plots show the proportion of each subset, with each T cell cluster represented similarly between samples.

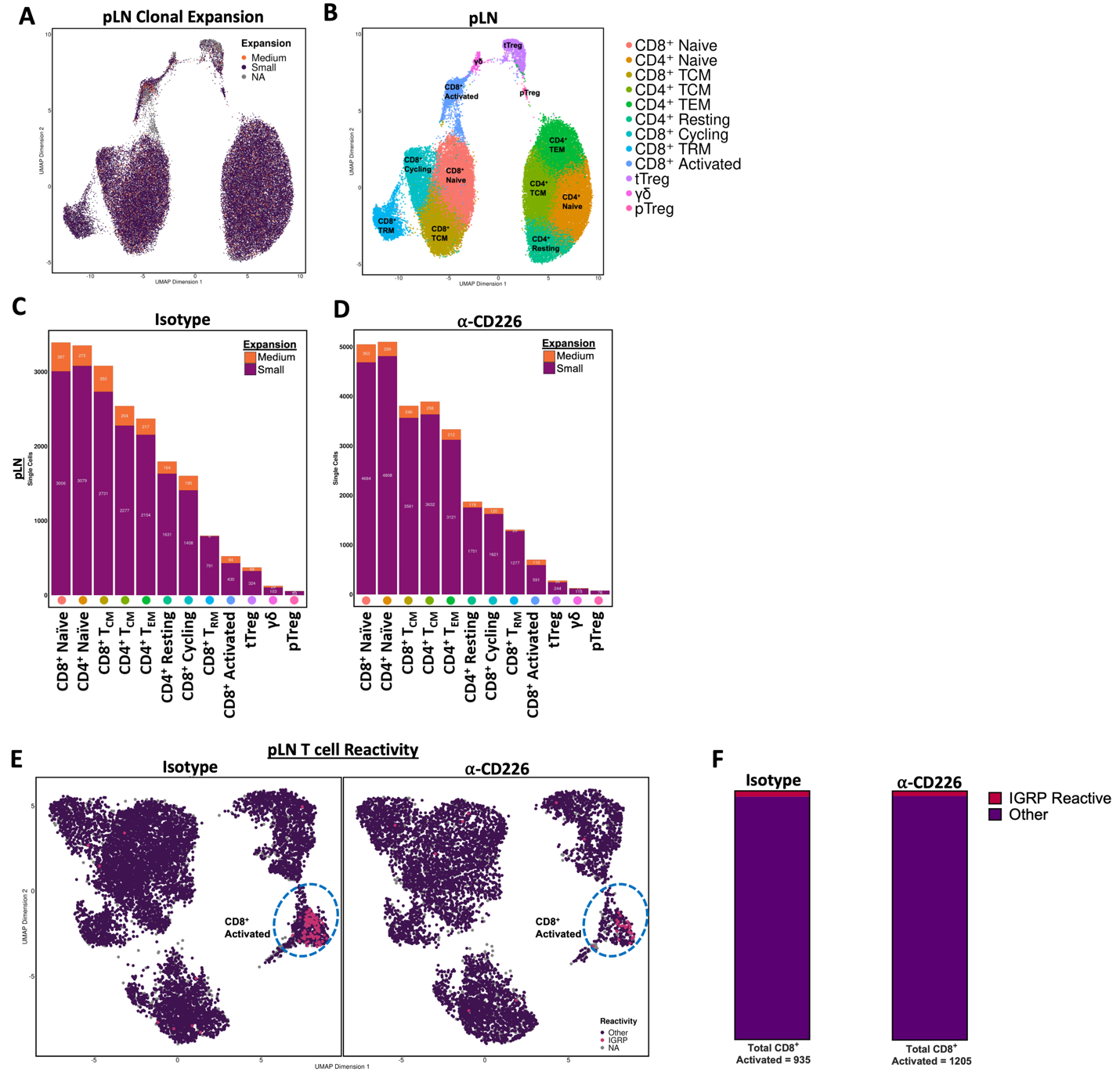
**Supplementary Figure 13. Anti-CD226 mAb Blockade Does Not Alter T Cell Clonal Expansion in the Pancreatic Lymph Node (pLN)**. To determine the impact of anti-CD226 mAb blockade on T cell clonal expansion in the pLN, TCR-seq data from each cell was merged with corresponding scRNA-seq data. (**A**) The degree of clonal expansion within each cluster was assessed by overlaying the proportion of each clonotype (Medium: 0.001 < X< 0.01 (orange), Small: 1e^-4^ < X < 0.001 (pink), NA: no CDR3 sequence for cell barcode (gray)) onto a Uniform Manifold Approximation and Projection (UMAP) of T cells isolated from NOD pLN, 5 weeks following treatment. (**B**) UMAP plots annotated with distinct T cell clustering for the pLN (17,617 cells). Stacked bar plots show the distribution of clonal expansion by cluster in mice treated with (**C**) isotype (**D**) or anti-CD226. (**E**) UMAP plots split by treatment condition show T cells containing CDR3 sequences within one amino-acid of previously published IGRP-reactive clones (Kasmani *et al.*), annotated as IGRP-reactive (pink), or as having another reactivity (purple). (**F**) Stacked bar plots show the frequency of IGRP-reactive clones within the CD8^+^ Activated T cell cluster between isotype or anti-CD226 treated mice. No significant P-values were identified using Fisher’s Exact Test between treatment conditions.
